# Cryo-EM Structure of Human ATAD2B Reveals a Hexameric Organization Contributes to ATPase Activity and Substrate Coordination

**DOI:** 10.64898/2026.04.02.716110

**Authors:** Kiera L. Malone, Eugene Y. D. Chua, James M. Lignos, Patricia M. Fagnant, Jill E. Macfarlane, Kathleen M. Trybus, Michael A. Cianfrocco, Karen C. Glass

## Abstract

ATPase family AAA+ domain-containing protein 2B (ATAD2B) is a poorly characterized member of the ATAD2-like protein family, which contains a unique combination of tandem AAA+ ATPase domains with a C-terminal bromodomain. In humans, ATAD2B is dysregulated in several disease states including cancer and respiratory disorders, yet despite its promise as a therapeutic target, little is known about its molecular function. Here, we report the first high-resolution cryo-EM structure of human ATAD2B at 3.0 Å, revealing a two-tiered hexameric assembly with a shallow spiral staircase architecture. Structural analysis uncovers conserved AAA+ ATPase features, including nucleotide coordination at inter-subunit interfaces, inter-subunit signaling (ISS) gate loops, and pore loops that engage a substrate within the central channel. Biochemical assays demonstrate that ATAD2B is an active enzyme with an ATP hydrolysis rate of 0.34 ATP/hexamer/sec. Furthermore, the integrity of the hexameric complex is stabilized through unique knob-hole interactions, a linker arm that extends between the AAA2 and bromodomain, and an N-terminal linker domain (LD). These findings establish ATAD2B as a functional AAA+ ATPase and provide mechanistic insight into its enzymatic activities, laying the foundation for understanding its role in chromatin-associated processes.

## Introduction

The human ATPase family AAA+ domain-containing 2B (ATAD2B) protein is a conserved epigenetic regulator implicated in transcriptional activation, chromatin organization, and DNA repair^1–4^. ATAD2B, and its ATAD2 paralog, are members of a unique family of human chromatin-associated factors distinguished by the presence of a bromodomain fused to tandem AAA+ ATPase domains^5,6^. Despite their shared ancestry, accumulating evidence indicates that ATAD2 and ATAD2B have divergent biological functions. ATAD2 is a well-established oncogene whose overexpression promotes transcriptional programs driving cell proliferation in multiple malignancies including breast, ovarian, and prostate cancers^7–10^. In contrast, ATAD2B displays a distinct expression profile, with emerging roles in neurological disorders, glioblastoma, and several respiratory diseases^1,11,12^. However, the molecular architecture of ATAD2B, and the mechanistic basis for its diverse disease-related associations are completely unknown.

Functional studies further highlight differences in the cellular roles of ATAD2 and ATAD2B. A systematic RNAi screen to identify human bromodomain-containing proteins involved in DNA double-strand break repair revealed that depletion ATAD2B impaired non-homologous end joining (NHEJ) DNA repair^4^. In comparison, siRNA knockdown of ATAD2 specifically reduces the efficiency of the homologous recombination DNA repair pathway^13^. Furthermore, ATAD2 is predominantly expressed in S-phase of the cell cycle where it associates with newly synthesized DNA to facilitate DNA replication^5^. Unlike the homologous recombination DNA repair mechanism, which uses the duplicate copy of DNA available only in the S and G2 cell cycle phases, the NHEJ DNA repair pathway operates throughout the cell cycle^14^. The association of ATAD2 with homologous recombination and ATAD2B with NHEJ DNA repair, suggests they have cell cycle specific roles. Differences between ATAD2 and ATAD2B also extend to how they engage with chromatin substrates via their bromodomains.

Biochemical profiling shows that the ATAD2B bromodomain recognizes a broader range of histone post-translational modifications (PTMs), including multiple combinations of PTMs on core and variant histones^2,15^. Notably, the ATAD2B bromodomain binds to acetylated histone H4 ligands that also contain nearby phosphorylation or methylation modifications, whereas the ATAD2 bromodomain does not^3^. ATAD2 preferentially recognizes newly synthesized H4 histones containing acetylated lysine 5 and 12 (H4K5acK12ac), which are found in S-phase during replication^5,6^. However, the ATAD2B bromodomain interacts with histone ligands that are common in euchromatin such as hyper-acetylated histone H4 (H4K5acK8acK12acK16ac), and the acetylated H2A.Z variant, which is typically added to regions of active gene transcription after cell division is completed^16–19^. These findings indicate that the two paralogs operate in separate chromatin regulatory pathways.

ATAD2 and ATAD2B are type II classical AAA+ ATPases, and are members of a larger ATAD2-like family of chromatin remodelers that are conserved from yeast to humans^1,20,21^.

While they share similar 2D architectures, including a flexible N-terminal domain, tandem AAA1 and AAA2 domains, a bromodomain, and a C-terminal domain (**Figure 1a**), significant sequence divergence and variations in their domain organization produce distinct functional behaviors. In yeast, the ATAD2-like homolog Abo1 uses ATP to load the H3-H4 histones onto DNA to prime nucleosome assembly. Another homolog known as Yta7 promotes nucleosome disassembly by threading the H3 tail into the central pore of its AAA+ hexameric ring^22–25^.

**Figure 1.**
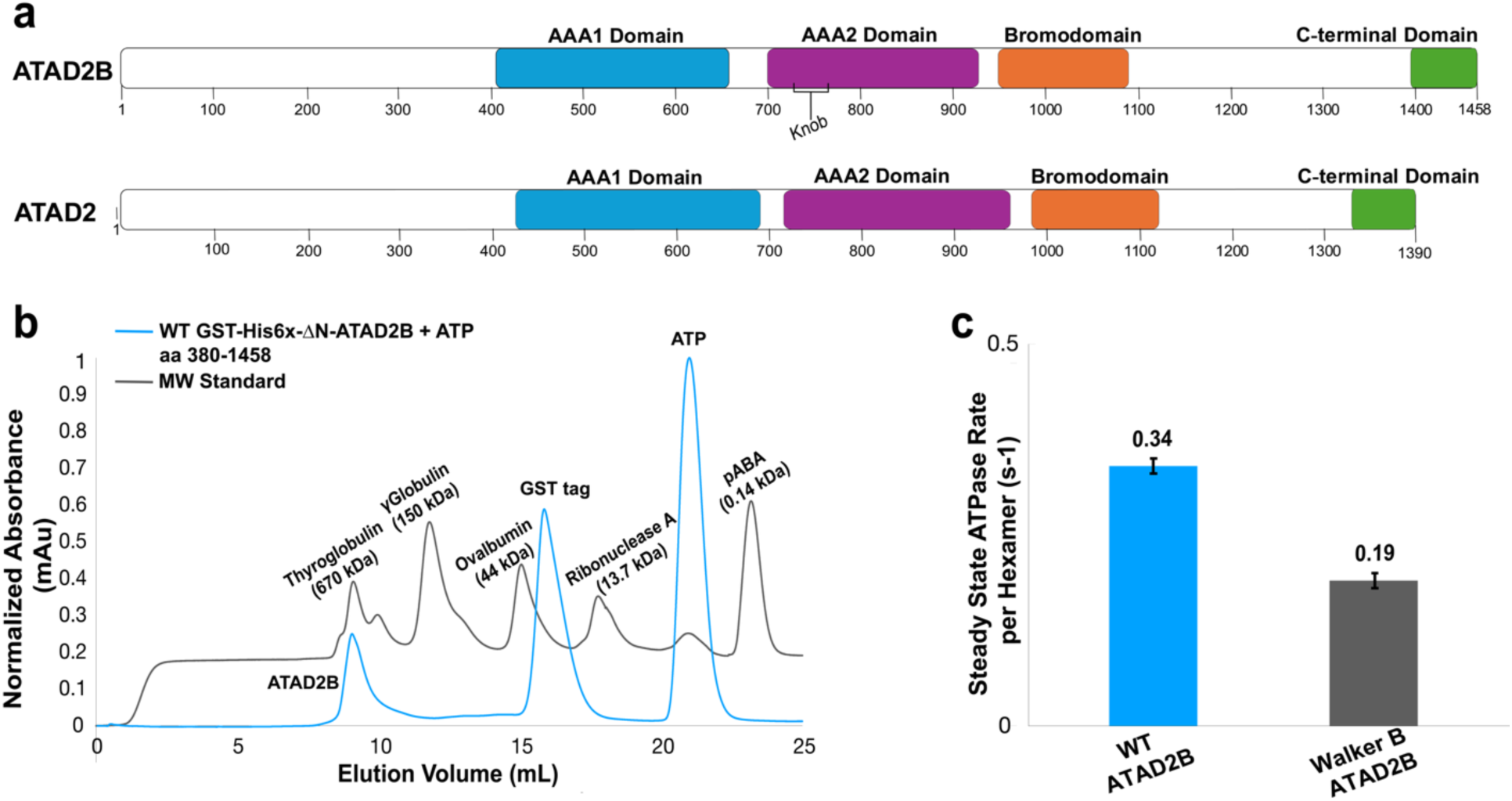
Human ATAD2B assembles into an active AAA+ ATPase. **a)** Domain organization of the full-length human ATAD2B (residues 1-1458) and ATAD2 proteins (residues 1-1390), with domains annotated. In ATAD2B the N-terminal domain runs from residues 1-406, the AAA1 covers residues 406-659, AAA2 covers residues 702-937, the knob region is residues 720-764, the linker arm runs from residues 917-954, the bromodomain covers residues 955-1085, and the C-terminal domain is residues 1398-1458. In ATAD2, the N-terminal domain consists of residues 1-431, the AAA1 covers residues 432-685, AAA2 covers residues 727-963, the knob region is residues 750-786, the linker arm runs from residues 944-980, the bromodomain covers residues 981-1110, and the C-terminal domain is residues 1330-1390. **b)** The size exclusion chromatography (SEC) trace from a Superdex 200 Increase 10/300 GL column for the wild-type (WT) GST-His6x-ΔN-ATAD2B (residues 380-1458) with 2 mM ATP in the buffer is shown in blue. The expected molecular weight for monomeric ATAD2B is 150 kDa. The SEC trace for the protein standard mix (Millipore Sigma, in grey) is overlayed with the ATAD2B trace and shows the first ATAD2B peak elutes with thyroglobulin (670 kDa), indicating ATAD2B forms a multimeric complex. The second peak is consistent with the GST-tag dimer (∼50 kDa) eluting near ovalbumin (44 kDa), and the final peak elutes prior to the pABA compound (MW 137 Da), which is consistent with free ATP in the buffer. **c)** The steady-state ATP hydrolysis rate for WT ATAD2B (blue bar) compared to the Walker B mutant ATAD2B (gray bar). Error bars represent the standard deviation of triplicate experiments done using the EnzChek^TM^ phosphate assay kit (Thermo Fisher Scientific).

Human ATAD2 (1390 aa) and ATAD2B (1458 aa) differ in the lengths of their linker regions connecting domains, and their N-terminal and C-terminal domains are not well-conserved^1^. Notably, the ATAD2 bromodomain maps to residues 981-1108, while the ATAD2B bromodomain occupies residues 953-1085, and their sequence differences produce distinct binding pocket geometries with different histone ligand binding preferences^3^. Structural studies on ATAD2, and the ATAD2-like homologs Abo1 and Yta7, reveal they assemble into hexameric complexes in which the tandem AAA+ domains form a two-tiered ring around a central pore^22,25,26^. The bromodomain regions of Abo1 and Yta7 form a third tier above the hexameric AAA1 ring, but their bromodomains do not recognize acetylated histones. Instead, their non-canonical bromodomains engage unmodified histone H3, which causes a pore opening conformational change^22,25^. The human ATAD2 AAA+ ATPase forms a similar hexameric complex that appears to adopt an autoinhibited state via a unique linker domain (LD)^26^. The ATP-hydrolysis activity of human ATAD2 was negligible, and the LD region inserted between the seam subunits is proposed to block catalysis. The absence of a structural model for human ATAD2B represents a major barrier to understanding how this unique AAA+ ATPase exhibits it distinct chromatin-binding preferences, unique cellular functions, and specific disease pathologies.

Here, we address this gap by determining the first high-resolution structure of human ATAD2B using single-particle cryo-electron microscopy (cryo-EM). The 3.0 Å cryo-EM reconstruction reveals that ATAD2B forms a two-tiered hexameric complex in which the AAA1 ring adopts an asymmetric spiral staircase conformation, while the AAA2 domain remains in a symmetrical planar ring. The hexamer is further stabilized through inter-subunit contacts that maintain oligomer integrity and contribute to the AAA+ ATPase and chromatin-binding activities of ATAD2B. Structural analysis identifies distinct nucleotide states across the AAA1 ring, with asymmetric coordination of ATP between subunits, and the seam subunits are bound to ADP. Conformational changes in the arginine fingers and inter-subunit signaling gate loops track these nucleotide states, indicating how the catalytic cycle of ATP-hydrolysis is propagated around the AAA1 ring. The catalytically dead AAA2 ring contributes important inter-subunit contacts that stabilize the hexamer formation via interlocking domains. Within the central pore, ATAD2B engages substrate density through conserved residues in the AAA1 pore loop regions that form a tryptophan staircase. Biochemical assays demonstrate that ATAD2B is an active AAA+ ATPase, suggesting it may couple the energy from ATP hydrolysis into mechanical force to translocate histone substrates into the central pore. These data establish the molecular architecture of ATAD2B, providing a structural framework for understanding how this highly significant but understudied chromatin regulator contributes to human disease.

## Results

### ATAD2B forms a stable hexameric AAA+ complex

Oligomerization into ring shaped complexes is a defining feature of AAA+ ATPases as it is essential for their function as molecular motors that couple the energy from ATP hydrolysis into mechanical force to translocate substrates into the central pore^21,27^. Previous structural studies on ATAD2-family proteins demonstrated that the yeast homologs Abo1 and Yta7 form a stable hexameric ring that is required for their histone chaperone and nucleosome disassembly activities^22,25^. Human ATAD2 also forms a hexameric complex, but displayed a more heterogenous range of oligomeric assemblies in both the presence and absence of ATP nucleotide^26^. The oligomerization behavior of human ATAD2B has not been examined. To determine whether ATAD2B forms a defined higher-order assembly, we first characterized its oligomeric state in solution.

To facilitate protein purification and biophysical analysis we adopted an approach successfully used for Abo1 and ATAD2 and removed the N-terminal region, which is predicted to be disordered in AlphaFold models^22,26^. The resulting ATAD2B construct contains an N-terminal GST-tag, a hexa-histidine (His6x)-tag, followed by residues 380-1458 of ATAD2B, which covers the two AAA+ ATPase domains, the bromodomain, and the C-terminal domain (**Figure 1a and Supplementary Figure 1**). Unless otherwise stated, this N-terminally truncated protein (GST-His6x-ΔN-ATAD2B, res 380-1458) is referred to as ATAD2B.

Recombinant ATAD2B was expressed using the bac-to-bac baculovirus expression system and purified by GST affinity chromatography followed by size-exclusion chromatography. In the presence of ATP, wild-type ATAD2B elutes as a well-defined oligomeric species in the first peak near thyroglobulin (**Figure 1b**, blue trace). To further evaluate the stability of this oligomer, we purified an ATP-hydrolysis defective Walker B mutant using the same protocol. In many AAA+ ATPases, Walker B mutations stabilize the ATP-bound state and promote formation of stable hexameric complexes^28–30^. Consistent with this behavior, as shown in **Supplementary Figure 2a** the Walker B mutant forms a stable oligomer without ATP added to the buffer as its elution peaks (pink trace) overlap with the wild-type ATAD2B bound to ATP (blue trace). We also examined the ATAD2B oligomeric assembly the presence and absence of ATP using negative stain electron microscopy. The particles in the micrographs of both wild-type and the Walker B mutant ATAD2B are the same size and form a ring-like shape, suggesting they form stable hexamers in the absence and presence of ATP (**Supplementary Figure 2b**). Compared with previously characterized homologs, wild-type ATAD2B appears more stable than human ATAD2, which displays a broad, heterogenous oligomeric distribution regardless of nucleotide state, but less stable than the Abo1 homolog in yeast, which forms highly stable hexamers independent of ATP^22,26^. Together, these results suggest that human ATAD2B forms a hexameric AAA+ ATPase.

### ATAD2B exhibits robust AAA+ ATPase activity

In AAA+ ATPase family proteins ATP hydrolysis is coupled to conformational changes around the ring and provides energy for substrate translocation^31^. In the ATAD2-like protein family, ATPase activity is required for histone loading or disassembly by the yeast homologs Abo1 and Yta7. In humans, deletion of the AAA1 domain prevents oligomerization of the ATAD2 complex and blocks its ability to interact with acetylated histone H4^32^. Although the ATP hydrolysis rate for human ATAD2 was found to be extremely slow in comparison to the Abo1 and Yta7 yeast homologs, ATP binding appears to play an important role in its function. Introduction of a Walker A mutation in human ATAD2, which prevents nucleotide coordination in other ATPases, was shown to inhibit oligomerization, binding to acetylated histone ligands, and reduced DNA replication^5,32,33^. We therefore sought to determine whether ATAD2B functions as an active AAA+ ATPase.

To measure ATP hydrolysis by ATAD2B, we employed an ATPase assay with the purified wild-type ATAD2B protein. In the presence of ATP, wild-type ATAD2B displays a steady-state ATP hydrolysis rate of 0.34 ATP per hexamer per sec (**Figure 1c**). This activity was reproducible across independent experiments and under *in vitro* buffer conditions used for biochemical and structural analysis of the ATAD2B complex. In contrast, the ATAD2B Walker B mutant (E506Q), which preserves ATP binding but disrupts catalytic priming of water for hydrolysis, showed a substantially reduced ATPase rate (**Figure 1c**), consistent with inhibition of ATP hydrolysis.

The ATP hydrolysis rate measured for ATAD2B is markedly higher than the rate previously reported for ATAD2, whose activity was close to the lower limit of detection in a comparable ATPase assay^26^. The ATPase activity of ATAD2B is within a similar range reported for the yeast homologs Abo1 and Yta7 at 0.83 and 0.43 ATP/hexamer/sec, respectively (**Supplementary Figure 2c**)^22,25^. Together these results demonstrate that ATAD2B is an enzymatically active AAA+ ATPase, similar to the ATAD2-like yeast homologs involved in histone chaperone and nucleosome disassembly functions.

### Cryo-EM reconstruction of human ATAD2B reveals an asymmetric hexameric assembly

High-resolution cryo-EM structures of the ATAD2-like family AAA+ ATPases Abo1, Yta7, and ATAD2 demonstrate that they share a conserved hexameric organization, yet adopt distinct conformations that correlate with differences in nucleotide bound states and chromatin-associated functions^22,25,26^. Since ATAD2B is an active AAA+ ATPase enzyme, we sought to determine its structure to define how nucleotide binding and oligomeric architecture are organized within this paralog. To stabilize ATAD2B in a nucleotide-bound state, we used the ATP-hydrolysis deficient Walker B mutant (E506Q), an approach previously shown to capture AAA+ ATPases in defined conformations^34^. To further promote hexamer stabilization and nucleotide occupancy, the protein was incubated with a slowly hydrolyzable ATP analog (ATPγS) and an acetylated histone H4K5acK12ac peptide (res 1-24), approximating a physiologically relevant ligand environment. Using this strategy, we determined a 3.0 Å cryo-EM reconstruction of human ATAD2B Walker B (E506Q) in the presence of ATPγS and H4K5acK12ac (**Figure 2, Supplementary Table 1**, **and Supplementary Figures 3 and 4)**.

**Figure 2.**
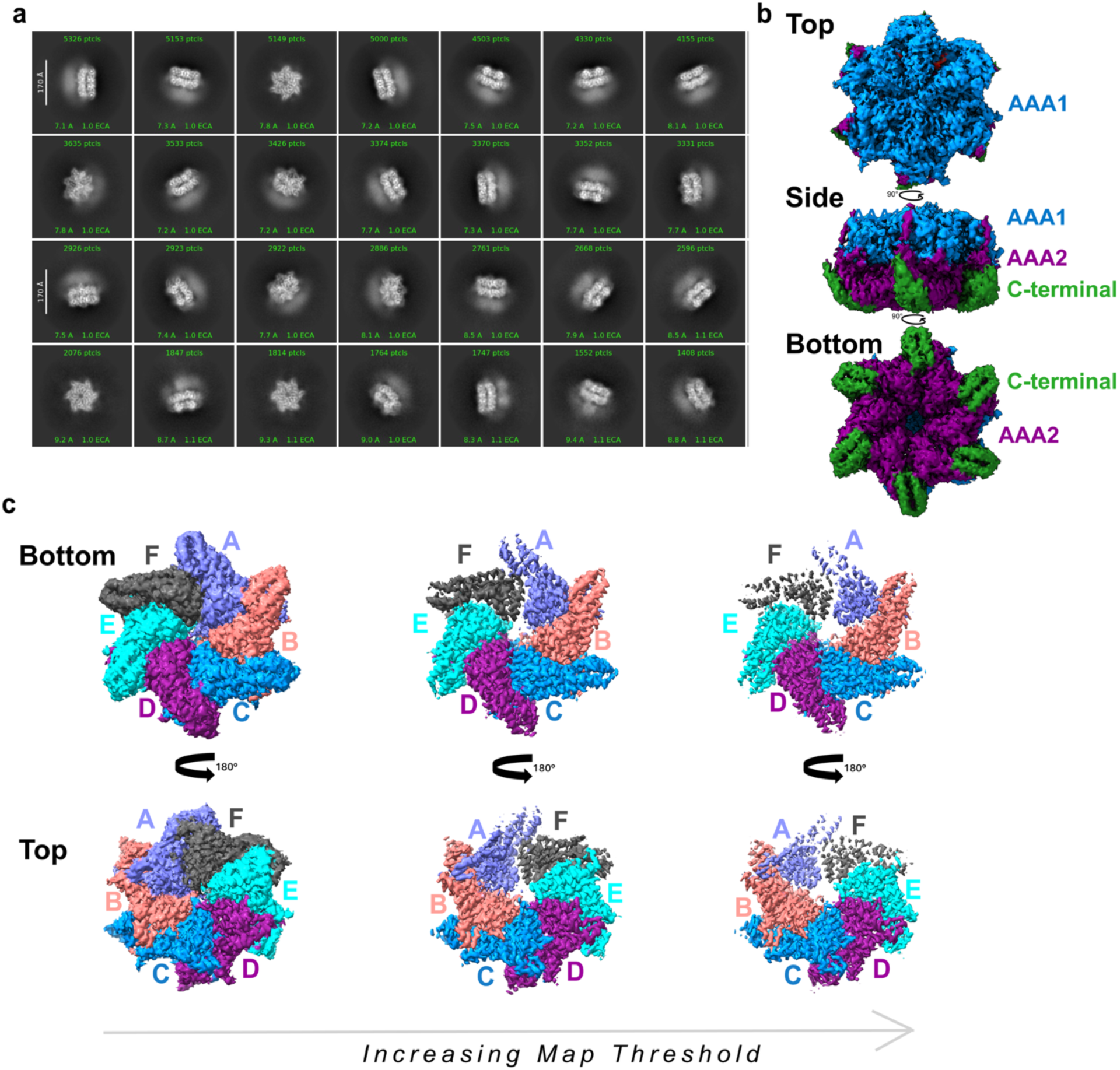
Cryo-EM structure of human ATAD2B reveals an asymmetric hexameric assembly. **a)** Selected 2D class averages of human ATAD2B Walker B (residues 380-1458, E506Q) with ATP*γ*S and H4K5acK12ac showing it forms a ring shaped hexamer. Side views appear to have three layers, one with diffuse density. These classes are representative of particles used in the final map reconstruction**. b)** Final cryo-EM reconstruction of the human ATAD2B Walker B complex (residues 380-1458, E506Q) with ATP*γ*S and H4K5acK12ac, shown from bottom, side, and top views. Ordered density is observed for the AAA1, AAA2, and C-terminal domains, whereas density corresponding to the bromodomain region is not resolved. The map is colored by domain as in Fig. 1a with AAA1 in blue, AAA2 in plum, and the C-terminal domain in clover. Density is displayed at a contour level of 0.02. **c)** Top and bottom views of the ATAD2B cryo-EM reconstruction colored by individual subunits within the hexameric assembly. Subunits A-F are labeled and colored as follows: Subunit A, orchid; subunit B, salmon; subunit C, aqua; subunit D, plum; subunit, turquoise; subunit F, iron. The bottom and top views are shown at increasing map contour thresholds (left to right: 0.02, 0.04, 0.06) to illustrate differences in local density quality and asymmetry in the subunits. Subunits A and F exhibit weaker density consistent with increased conformational flexibility. The curved arrows indicate rotation between views.

The 2D class averages of cryo-EM images of the ATAD2B particles show a ring shaped hexamer with views that suggest three vertical layers, one of which has diffused density (**Figure 2a).** The final 3D reconstruction shows a two-tiered hexamer where the AAA1 ATPase domains form a ring layer above the AAA2 ring, and the helical C-terminal domains form an extension associated with the lower part of the AAA2 domains (**Figure 2b**). This organization places ATAD2B in the type II classical AAA+ ATPase family, which assemble into two stacked hexameric rings via contacts between their AAA+ domains^35–37^. While the 2D classes show diffuse density above the AAA1 domain that is consistent with the location of the bromodomain observed in the Abo1 and Yta7 structures^22,25^, it is too weak for further analysis or model building (see blurry region in the 2D class averages) (**Figure 2a,b**). This is likely due to inherent flexibility in the bromodomain region, which is positioned at the top of the AAA+ stack via a long unstructured region that connects to the C-terminal bromodomain.

Coloring the cryo-EM map by individual subunits revealed pronounce asymmetry within the hexamer (**Figure 2c**). While most of the subunits have well resolved density across the AAA1 and AAA2 rings, subunits A and F exhibit notably weaker density, which becomes progressively attenuated at higher contour thresholds. These two subunits meet at a flexible seam interface, which is a conserved feature among AAA+ ATPases that catalyze ATP hydrolysis around the ring to engage substrate in the central pore^31,34,36,38,39^. Together, these data demonstrate that ATAD2B assembles into an asymmetric hexameric complex with AAA1 and AAA2 forming a well ordered two-tiered ring, featuring a central pore and increased flexibility at the seam subunits.

### The AAA1 ring forms a shallow spiral above the planar AAA2 hexameric ring

Having established the overall architecture of the ATAD2B hexamer, we next examined how individual subunits are arranged relative to each other to determine how this organization gives rise to the asymmetric assembly. ATAD2B contains two AAA+ domains in tandem, which classifies it among the type II AAA+ ATPases in the functionally diverse classical clade^21^. The AAA+ domains contain canonical structural features including two subdomains consisting of a nucleotide binding domain (NBD) followed by a smaller helical bundle domain (HBD)^30^.

The two-tiered ring formed by the AAA1 and AAA2 domains in ATAD2B is relatively planar, but contains a shallow rise (**Figure 3a,b**). Close inspection reveals that the AAA2 ring is nearly planar and symmetric, whereas the AAA1 ring adopts a shallow spiral staircase configuration. In this arrangement, the AAA1 domain of subunit F occupies the highest position in the ring, and the AAA1 ring descends downwards is a clockwise direction through subunits E, D, C, and B, with A forming the lowest step of the staircase (**Figure 3a, bottom right panel)**. The top and bottom subunits in the spiral staircase (subunits F and A), comprise the seam interface, consistent with their weaker density and lower resolution in the cryo-EM map due to increased flexibility (**Figure 2c and Supplementary Figure 5**).

**Figure 3.**
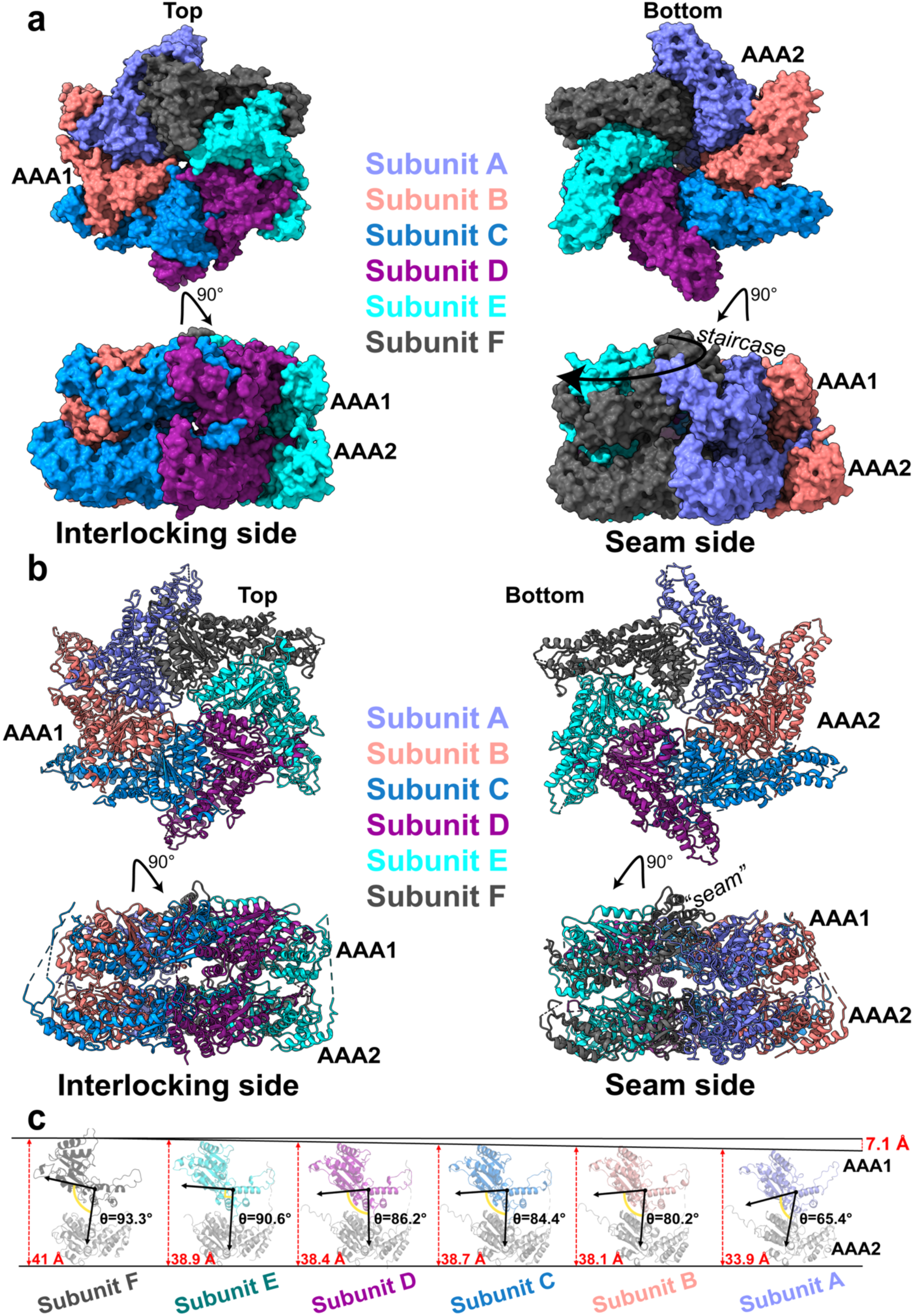
A shallow AAA1 spiral staircase underlies ATAD2B hexamer organization. **a)** Surface representations of the ATAD2B Walker B hexamer colored by subunit with subunit A in slate, subunit B in salmon, subunit C in marine, subunit D in purple, subunit E in cyan, and subunit F in gray. Top and bottom views of the complex are shown, along with side views of the complex rotated by 90° to highlight the more rigid ‘interlocking’ subunit interface, and the flexible ‘seam’ interface. The seam is formed at the junction between subunits A and F, and the shallow AAA1 staircase is indicated in the seam view. **b)** Cartoon representations of the ATAD2B Walker B hexamer shown in the same orientations as in (a), illustrating secondary structure features within the complex. The relative positions of the AAA1 (upper) and AAA2 (lower) rings are indicates, and the seam between subunits A and F is labeled. **c)** Comparative analysis of the subunit positions along the spiral staircase. Individual subunits were aligned by their AAA2 domains (shown in light grey) and arranged from the highest (subunit F) to lowest (subunit A) moving down the AAA1 staircase. The AAA1 domains are shown on top and colored by subunit. The angle between the AAA1 and AAA2 domains was measured using residues 548, 611, and 870, and is indicated by black arrows and a yellow angle (θ). The vertical offset between the nucleotide binding domains of AAA1 and AAA2 (relative distance between residues 570 and 895 in each subunit) is shown by red dashed lines with the distance indicated. The cumulative decrease in height between the F and A subunits around the ring is 7.1 Å, as indicated by the red dashed line.

To quantify the geometry of this spiral arrangement, the individual subunits were structurally aligned by their AAA2 domains, which are highly superimposable with root mean square deviation (RMSD ∼0.4-0.7 Å), enabling a direct comparison of the different positions adopted by the AAA1 domains (**Figure 3c and Supplementary Figure 6).** This analysis reveals the cumulative vertical offset between the highest AAA1 domain (subunit F) and the lowest AAA1 domain (subunit A) is 7.1 Å. Within the AAA1 domain of each subunit, the orientation of the NBD to the HBD remains relatively constant in the non-seam subunits B-E (1-2 degree differences, shown in **Supplementary Figure 7**). However, the angle between the large NBD and small HBD changes by 7 degrees between subunits B to A, and by 11 degrees from subunit F (118 °) to subunit A (129 °) at the seam. As shown in **Figure 3C** the angle between the AAA1 and AAA2 domains of each subunit progressively decreases by approximately 4 ° between successive subunits from F to B, followed by a more pronounced 15 ° decrease between subunits B to A. Moving clockwise around the ring from subunit A to F there is a large 28 degree change in the relative orientation of the AAA1 to the AAA2 domain in each subunit.

These measurements indicate that the shallow AAA1 spiral arises from conformational changes in the AAA1 domains that generated by rigid body rotations between the NBD and the HBD, and progressive changes in the orientation of the AAA1 domain relative to the AAA2 domain. Thus, a gradual downward movement of the AAA1 domain around the hexameric ring culminates at the seam, providing a structural basis for the observed asymmetry and localized flexibility. Together, these data establish that human ATAD2B adopts a shallow spiral staircase through conformational changes in the AAA1 domains, and their position above the planar AAA2 ring.

### Asymmetric ATP binding and sequential hydrolysis in the ATAD2B AAA1 ring

The shallow spiral staircase formed by the AAA1 domains and the localized flexibility at the A-F seam subunits (**Figure 3**), is characteristic of AAA+ ATPases that hydrolyze ATP sequentially around the ring ^31,34,40^. We therefore analyzed nucleotide identity, local coordination, and inter-subunit catalytic contacts across the AAA1 ring to determine if ATAD2B displays additional characteristics of a rotary ATP hydrolysis mechanism.

Inspection of the AAA1 domain architecture confirms it adopts the conserved AAA+ fold consisting of a larger Rossmann-like nucleotide binding domain (NBD) coupled to a smaller helical bundle domain (HBD) (**Figure 4a**). In the ATAD2B Walker B (E506Q) cryo-EM reconstruction, density corresponding to the expected nucleotide binding location is present at the interface of every subunit (**Figure 4b-h**). Notably, five of the AAA1 subunits (B-F) are occupied by ATPγS, while subunit A at the bottom of the spiral contains ADP. This asymmetric nucleotide state around the ring is consistent with cryo-EM structures of other ATAD2-like family members, and is indicative of the catalytic state of subunits in the AAA+ ATPase complex^22,25,26,30,31^.

**Figure 4.**
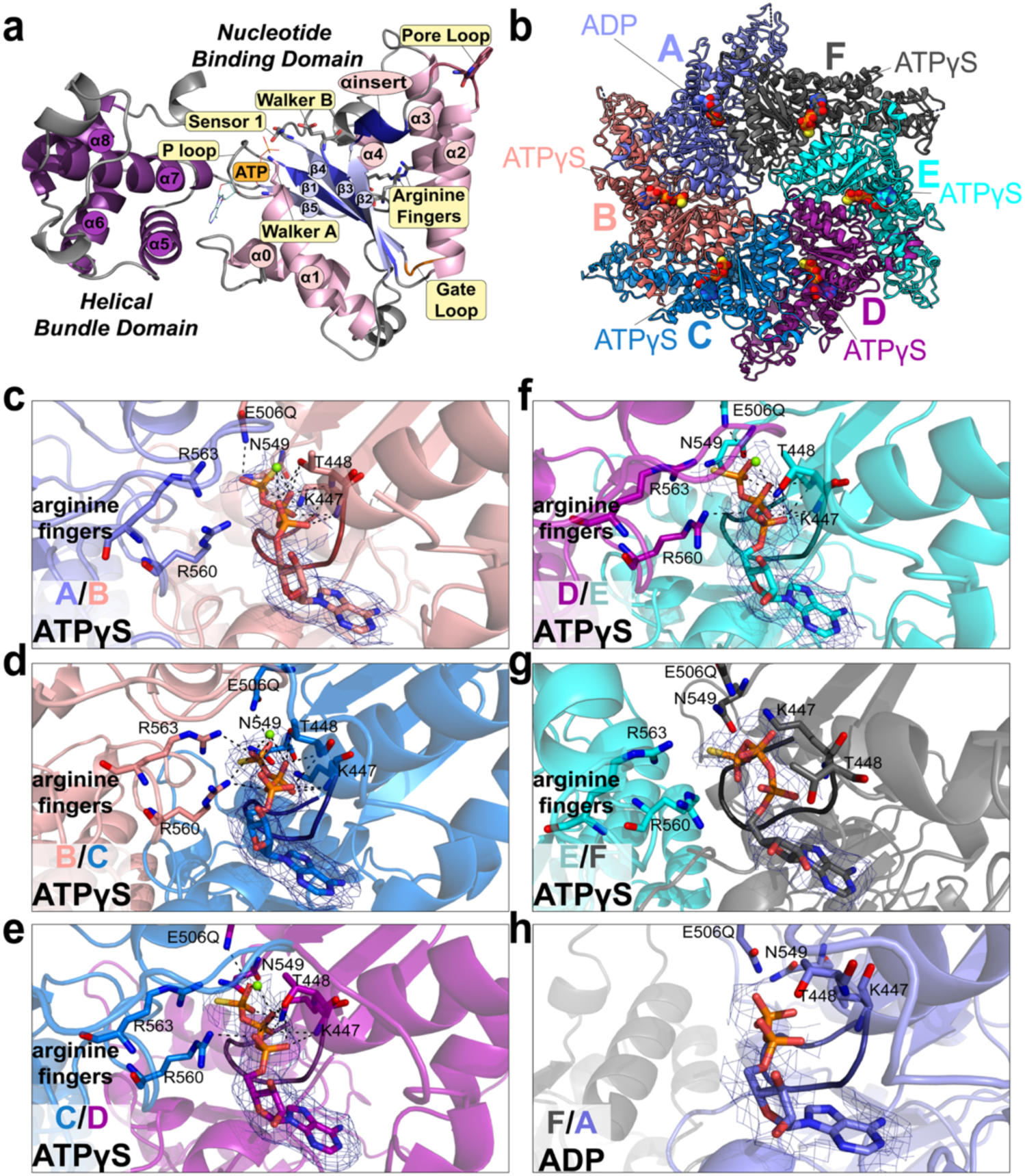
Nucleotide occupancy and coordination across the ATAD2B AAA1 ring. **a)** Schematic representation of the AAA1 domain from the AlphaFold model of ATAD2B (AF-Q9ULI0-F1-v4) highlighting conserved structural elements of the AAA+ fold, including the nucleotide binding domain (NBD; Rossmann-like large subdomain), the smaller helical bundle domain (HBD), and connecting linker regions. Conserved residues involved in nucleotide coordination and inter-subunit communication are shown as sticks and labeled. Nucleotide binding and hydrolysis residues include the Walker A P-loop residue Lys447, the Walker B residue E506Q, Sensor 1 Asn549, and the arginine fingers (Arg560 and Arg563). The gate-loop functions as an inter-subunit signaling motif (residues 534-541), and the conserved pore loop 1 residue W479 that engages substrates in the pore. The ATP binding position is also indicated. **b)** cartoon representation of the ATAD2B Walker B hexameric complex colored by subunit, showing the locations of bound nucleotide ligands. **c-h)** Close-up views of the AAA1 domain nucleotide-binding pockets found at the interface of adjacent subunits A-F. Each nucleotide is oriented to highlight residues from the cis-and trans-acting AAA1 domains in each subunit. ATPψS (in AAA1 of subunits B-F) or ADP (in AAA1 of subunit A) are shown as sticks, together with surrounding residues that form hydrophobic or polar contacts to the nucleotides. Cryo-EM density surrounding each nucleotide is shown as a blue mesh (contoured at σ=5.0 in PyMOL). In each binding pocket the conserved cis-acting residues include the Walker A P-loop residues Lys447 and Thr448, the Walker B residue E506Q, and Sensor 1 Asn549. Trans-acting residues in the AAA1 domain of the clockwise neighboring subunit include the arginine fingers (Arg560 and 563). Mg^2+^ ions are shown as green spheres where observed, and polar contacts are indicated by dashed lines. Additional views of density in the AAA1 nucleotide binding sites are shown in **Supplementary Figure 8**.

Across the AAA1 ring, nucleotide binding occurs in a cleft between the NBD and HBD (**Figure 4c-g**). While clear density is observed for each nucleotide, the density for the nucleotide coordinated between the seam subunits A and F is weaker, consistent with the flexibility of this region (**Supplementary Figure 8**, **compare a-d to e-f**)^40^. The adenine base is positioned between the N-terminal helices of the NBD and the four alpha-helical bundle in the HBD, while the phosphate groups are coordinated by residues in the subunit interface. These include the Walker A P-loop residues Lys447 and Thr448, the Walker B residue (E506Q), and sensor 1 residue Asn549 from the same subunit coordinating the adenine group. In addition, trans-acting residues from the clockwise neighboring subunit extend into the nucleotide binding pocket to stabilize ligand binding. This includes the arginine fingers (Arg560 and Arg 563), and the inter-subunit signaling (ISS) “gate loop” motif (residues 534-541)(**Figure 4b-e**). Density for a coordinating Mg^2+^ ion is evident in the ATPγS sites of subunits B-E, but is absent in the seam subunits (**Figure 4c-h**).

Comparison of individual nucleotide binding pockets reveals systematic changes in the inter-subunit coordination of nucleotides around the AAA1 ring. In subunits B, C, and D, the arginine fingers reach from the neighboring subunit to make hydrogen bond contacts with the ATPγS phosphate groups of the nucleotide bound in the adjacent subunit. This observation is consistent with their role in inter-subunit communication by sensing the nucleotide-bound state^21,31^. The hydrogen bonds between the arginine finger residues and the nucleotide bound in the next subunit are lost in the E/F and F/A interfaces, where increased inter-subunit spacing correlates with reduced arginine finger engagement and diminished nucleotide density (**Figure 4g,h and Supplementary Figure 8**). In subunit A, which contains ADP, the arginine fingers from the clockwise subunit F have retracted away from the nucleotide binding pocket, indicating a post ATP hydrolysis configuration. Reduced in trans contacts are also observed at the A/B interface, consistent with subunit B being poised to undergo the next round of ATP hydrolysis as the cycle progresses around the ring^41^.

Unlike in the AAA1 domains, no density corresponding to bound nucleotide is observed in the AAA2 domain of any subunit. In the ATAD2-like family, the second AAA+ domain (AAA2) is catalytically inactive because it is missing the conserved Walker A motif and arginine finger residues that coordinate ATP, as well as the Walker B motif that contributes to ATP hydrolysis (**Supplementary Figure 6**)^22,25,26^. Despite lacking catalytic activity, the AAA2 ring of ATAD2B is well ordered and planar, which provides a rigid structural base to support ATP hydrolysis in the asymmetric AAA1 spiral ring.

Collectively, the asymmetric distribution of ATPγS/ADP nucleotides, the gradual loss of arginine finger and Mg^2+^ ion coordination of nucleotides in the A/F seam subunits, and the absence of nucleotide binding in the planar AAA2 ring support a model in which ATP hydrolysis proceed sequentially around the AAA1 ring, while AAA2 functions as a non-catalytic structural scaffold. The divided roles of the AAA1 and AAA2 domains in each subunit explains how ATAD2B couples asymmetric AAA1 conformational changes in a stable hexameric complex throughout its ATPase cycle.

### Gate loop ISS motifs adopt seam dependent conformations in the ATAD2B hexamer

The asymmetric AAA1 ring observed in ATAD2B (**Figures 3 and 4**) is characteristic of the ATP hydrolysis cycle and suggests that additional inter-subunit elements beyond the arginine fingers may participate in sensing and propagating nucleotide-dependent conformational changes^34^. In AAA+ ATPases, the inter-subunit signaling (ISS) motifs (also known as gate loops) have been shown to coordinate communication across neighboring subunits^31^. We therefore investigated the conformation and position of the gate loops in ATAD2B to determine how they relate to nucleotide binding and seam formation within the hexamer.

In the ATAD2B Walker B structure, the gate loops extend between adjacent subunits and partially cover the nucleotide binding pocket of the clockwise neighboring subunit (**Figure 5a and Supplementary Figure 9**). Within the hexamer two distinct gate loop conformations exist, where subunits with ‘open’ and ‘closed’ positions are grouped around the ring (**Figure 5a**).

**Figure 5.**
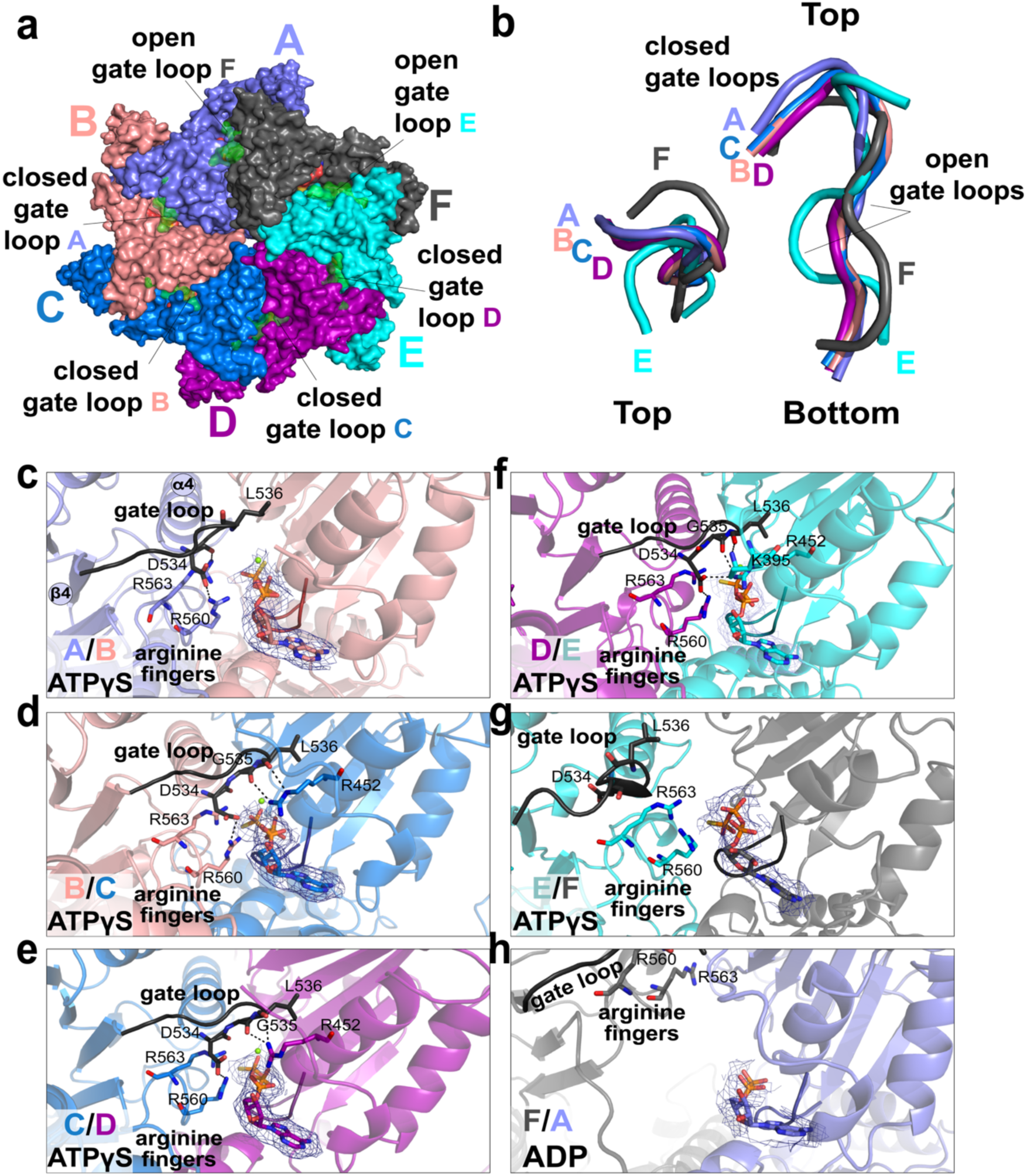
Gate-loop conformational changes are associated with nucleotide coordination. **a)** Surface representation of the ATAD2B Walker B hexamer colored by subunit, with the AAA1 gate loops (residues 534-541) highlighted in green and shown at 50% transparency. Subunits A-D adopt closed gate loop conformations, while subunits E and F adopt open conformations. **b)** Images representing segments of the gate loops from all six subunits aligned to subunit C, shown in top and side views. The overlayed gate loops highlight the conformational differences between subunits with closed gate loops (subunits A-D) versus subunit with open gate loops (subunits F and E). **c-h)** Close up views of the AAA1 subunit interfaces A/B (b), B/C (c), C/D (d), D/E (f), and F/A (g) in the ATAD2B Walker B structure. Protein subunits are shown as cartoons, with residues contacting the nucleotide ligands drawn as sticks, and the gate loop regions are shown in black. The bound nucleotides ATPγS (subunits B-F) or ADP (subunit A) are depicted as sticks. Cryo-EM density corresponding to each nucleotide is shown as blue mesh (contoured at σ=5.0 in PyMOL). The arginine finger residues are labeled, polar contacts are indicated by dashed lines, and Mg^2+^ ions are shown as green spheres where observed. Additional views of gate-loop density are shown in **Supplementary Figure 9**.

These gate loops are located between the α3-helix and β4-strand of the AAA1 domain (**Figure 5b**), and correspond to the conserved ISS motif described in other AAA+ ATPases^31,34^. In ATAD2B, the gate loop/ISS motif comprises residues Asp534, Gly535, and Leu536, and the third residue in this conserved motif is typically a hydrophobic aromatic residue, or Leu, Met, or Val (**Supplementary Figure 1**)^31^. As shown in **Figure 5c-f**, subunits A-D adopt a closed gate loop conformation, in which the ISS motif extends into the adjacent subunit over the nucleotide binding pocket. In contrast, subunits E and F contain gate loops in the open conformation (**Figure 5f,g**). Alignment of individual gate loops relative to subunit C shows that the closed conformations overlap closely, while the open conformations from subunits E and F are displaced and had distinct backbone trajectories (**Figure 5a, middle and right**), indicating a reproducible and position dependent structural rearrangement.

In subunit interfaces A/B, B/C, C/D, and D/E, the closed gate loops engage the adjacent nucleotide binding pocket through a combination of hydrophobic and polar contacts (**Figure 5b-e and Supplementary Figure 10**). Leu536 inserts into a hydrophobic cleft formed by beta-strands of the neighboring NBD, while the backbone carboxyl of Gly535 and Leu536 make polar contacts with Arg542 in the adjacent subunit for efficient gate loop closure. Asp534 stabilizes the arginine fingers in cis through additional hydrogen bonds (**Figure 5b-e**).

In contrast, at the interfaces of subunits E/F and F/A, the gate loops are retracted away from the nucleotide binding pocket (**Figure 5f,g and Supplementary Figure 10**). The gate loop for subunit E is lifted upward and no longer makes hydrogen bond contacts R452 or the arginine finger residues, and the gate loop of subunit F has moved even further away from the nucleotide binding pocket of the adjacent subunit. Notably, the subunit interfaces with closed loops (A/B, B/C, C/D, and D/E) contain density features that are consistent with ATPγS, while the E/F and F/A interfaces have reduced nucleotide density. While the nucleotide density at the E/F interface is most consistent with ATPγS, the EM density observed at the F/A interface is smaller, and more consistent with ADP being bound at the seam (**Figure 5g and Supplementary Figure 9**).

Together, these data demonstrate that the ATAD2B gate loops/ISS motifs participate in inter-subunit communication and sensing of the nucleotide bound state. Closed gate loops stabilize the nucleotide binding interface, whereas the open gate loops may facilitate a path for nucleotide exchange. Differences in the gate loops of subunits surrounding the seam interface provides a structural basis by which conformational switches in ATAD2B are used to sense nucleotide states and transmit inter-subunit signals around the AAA1 ring during its ATPase cycle.

### A knob-hole interaction stabilizes the ATAD2B hexameric assembly

AAA+ ATPases frequently acquire functional diversity through extensions to their sequences and insertion of functional domains, which can stabilize oligomeric assemblies or modulate their inter-subunit contacts^21,30,31,42,43^. In the ATAD2-like family, prior cryo-EM studies of Abo1 and Yta7 identified a distinctive knob-hole architecture emanating from insertions in the AAA2 domain and an extended linker arm between AAA2 and the bromodomain, respectively, that together stabilize the hexameric assembly^22,25^. Given the asymmetric organization and seam flexibility observed in ATAD2B (**Figure 3-5**), we examined if corresponding stabilizing features are present in the ATAD2B structure and how they compare with related ATAD2-like family members.

Inspection of the ATAD2B Walker B cryo-EM map reveals ordered density beyond the β1 strand of the AAA2 domain (residues 750-764), corresponding to a knob-like insertion that is not part of the canonical AAA+ fold (**Figure 6a, and Supplementary Figure 1**). Although this region is predicted to be a disordered loop in the AlphaFold models of ATAD2B (AF-Q9ULI0-F1-v4)(**Supplementary Figure 11b**)^44^, partial density is observed in several subunits, allowing model building near residue Pro750. The knob projects counter-clockwise into the adjacent subunit and pack against the neighboring AAA1 domain, forming an interlocking contact at select interfaces (**Figure 6a and Supplementary Figure 11**).

**Figure 6.**
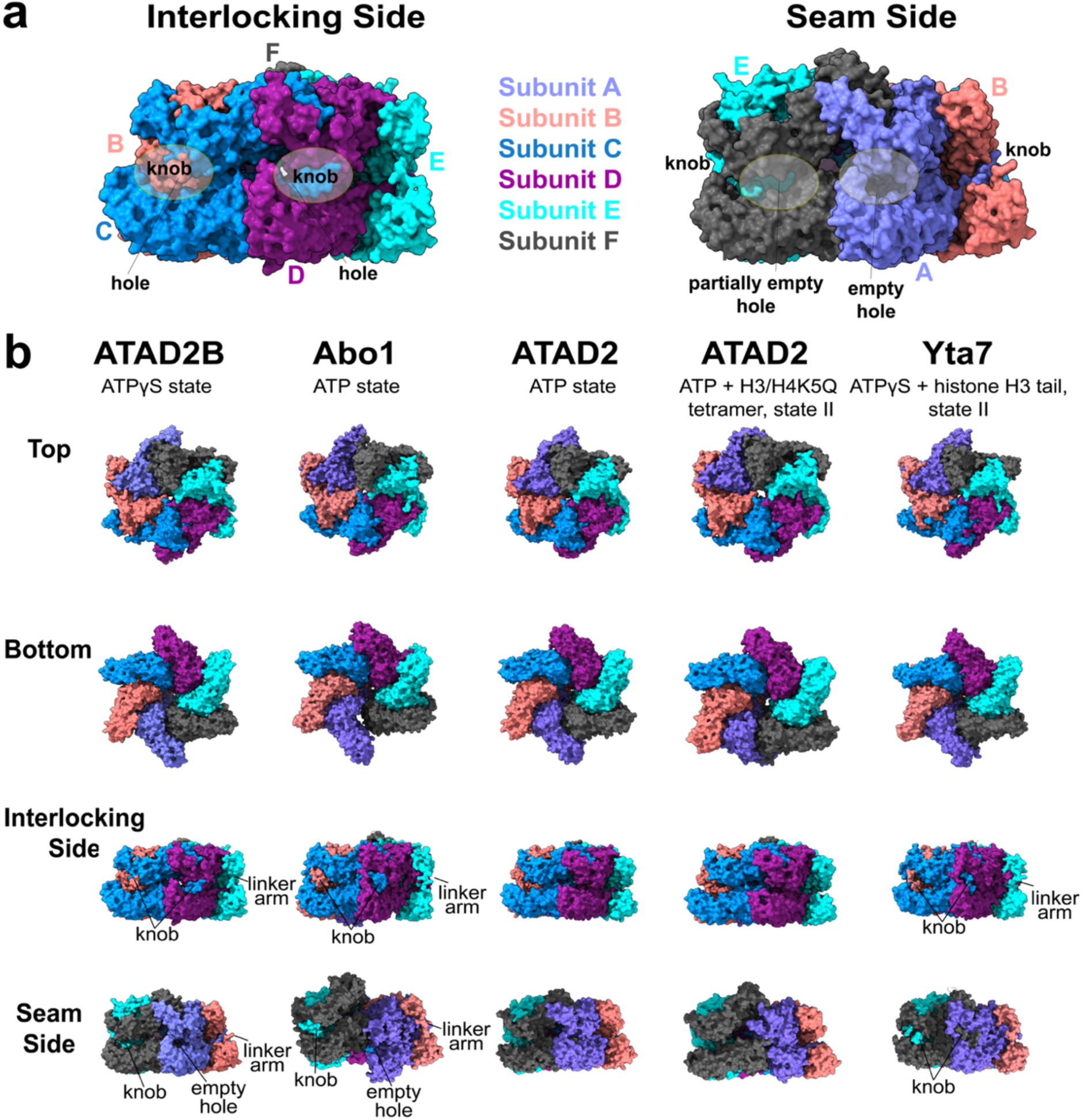
The knob-hole and linker arm architecture stabilizes the ATAD2B hexamer. **a)** Surface representations of the ATAD2B Walker B structure colored by subunit and shown from the interlocking subunit side (left) and the seam side (right). The AAA1 domain is oriented at the top and the AAA2 domain is on the bottom. Transparent gray circles indicate the positions of the ‘knob’ protrusions and corresponding ‘hole’ cavities at the inter-subunit interfaces. On the interlocking side, each knob inserts into a matching hole on the adjacent subunit, whereas on the seam side one hole is unoccupied (‘empty hole’), reflecting the absence of a stabilizing knob at this interface. **b)** Comparison of oligomeric architectures across the ATAD2-like family of AAA+ ATPases. Surface representations are shown for ATAD2B bound to ATPγS (this study), followed by Abo1 Walker B-ATP (PDB ID: 6jq0), ATAD2 Walker B-ATP (PDB ID: 8h3h), ATAD2 Walker B-ATP bound to the H3/H4K5Q histone tetramer (state II, PDB ID: 8juy), and Yta7 Walker B-ATPγS bound to a histone H3 peptide (state II, PDB ID: 7uqj). For each complex, top (AAA1), bottom (AAA2), interlocking side, and seam side views are shown. All structures were aligned the ATAD2B using Subunit C in ChimeraX^45^. The C-terminal knob-hole regions, and the linker arm connecting the AAA2 domain to the bromodomain are labeled.

The knob density is most clearly resolved at non-seam subunit interfaces (A/B, B/C, C/D, and D/E), where it inserts into a complementary hole on the adjacent subunit (**Figure 6a**). In contrast, density for the knob is weak or absent at the subunit interfaces moving clockwise from the seam (e.g. F/A and E/F). In particular, no knob extension is observed at the interface of the seam subunits F and A, which is similar to the ATP-bound structure of Abo1 where knob engagement is also absent at the seam^22^. Interestingly, we only observe density for the knob starting at residue Tyr759 in subunit E (rather than at residue 750 in A-D), which prevents the visualization of the knob counterclockwise extension from E into subunit F as with other subunits (**Figure 6**). These observations indicate that knob-hole interactions preferentially stabilize non-seam subunit interfaces with the ATAD2B hexameric assembly, enabling increased motion at the seam.

The hole that accommodates the knob is formed in between the HBD of AAA1 and the HBD of AAA2 along with the linker arm that runs up the side of the two-tiered ring (**Figure 6a and Supplementary Figure 11a**). The linker arm of ATAD2B originates from the end of the AAA2 and includes residues 920-937, which form the unstructured region between AAA2 and the bromodomain at the top of the hexamer (**Figure 2a**). Although continuous density for the full linker arm sequence is not observed, partial density consistent with this trajectory is visible in the non-seam subunits B-E, opposite the seam (**Figure 6a, left and 6b**). As with the knob, density for the linker arm region is weakest or absent in the seam subunits A and F.

Comparison of ATAD2-like structures highlights the similarities and differences in these stabilizing elements (**Figure 6b**). The cryo-EM maps of Abo1 and Yta7 display robust knob-hole engagement with show well resolved density to build the linker arm regions across most non-seam subunits^22,25^. ATAD2B shows partial ordering of the linker arm and knob structures, while ATAD2 lacks continuous density for the knob and linker arms, even the interlocking subunits^26^. These structural differences may be indicative of the relative stability of the hexameric assemblies observed among the ATAD2-like family members.

Collectively, these results demonstrate that ATAD2B contains a knob-hole and linker arm architecture that stabilizes the interlocking subunit interfaces within the hexameric complex, while these structural features are disordered in the more flexible seam subunits. The partial density of these features places ATAD2B between ATAD2 and the more stable Abo1 and Yta7 complexes, providing structural insight into how differences in oligomer stabilization contributes to functional diversification within the ATAD2-like family of AAA+ ATPases.

### An N-terminal linker domain occupies the hexameric seam

Multiple stabilizing features of the ATAD2B hexamer, including nucleotide coordination, gate loops, and the knob-hole interaction, have pronounced asymmetry near the seam subunits (**Figures 3-6**). In related ATAD2-family AAA+ ATPases, an N-terminal linker domain (LD) preceding the AAA1 module has been shown to localize at the seam and influence inter-subunit packing^22,25,26^. However, prior structures indicate that LD engagement varies across family members and conformational states. We therefore examined the ATAD2B cryo-EM map to determine if a comparable LD is present, and how its conformation relate to seam architecture and homologous ATAD2 structures.

The ATAD2B Walker B-ATPγS-H4K5acK8ac cryo-EM map reveals additional density at the N-terminus of the AAA1 domain in subunit A, corresponding to residues 389-398 (**Figure 7a-c and Supplementary Figure 12**). This density is absent for residues 389-392 in the analogous region of subunits B-F, indicating that ordering of the N-terminal LD region is restricted to the seam. In the interlocking subunits the shorter LD segment is overlayed on top of the subunit interface (**Figure 7b**), while the extended LD region of subunit A packs into the cleft between the AAA1 domains of subunits A and F (**Figure 7c**).

**Figure 7.**
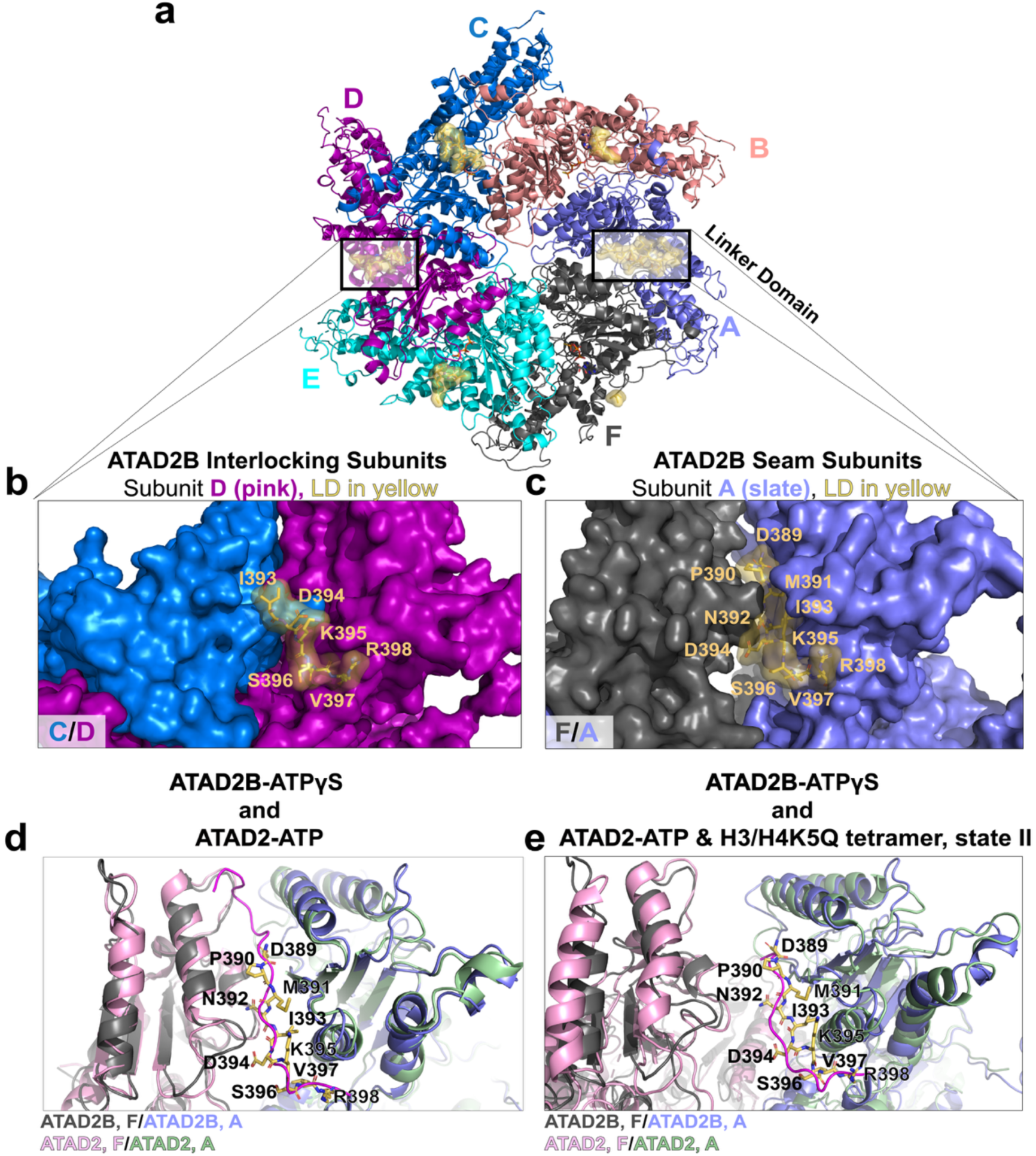
The ATAD2B linker domain adopts distinct conformations at interlocking and seam subunits. **a)** Top view of the ATAD2B Walker B hexamer is shown in cartoon and colored by subunit. The residues of the N-terminal linker domains are colored yellow and highlighted by boxed regions, with representative interlocking and seam interfaces indicated. **b)** Close-up surface view of the linker domain position at the interface of interlocking subunits C/D, where residues 393-397 from subunit D are well resolved. **c)** Close-up surface view of the linker domain at the interface of the seam subunits F/A, showing residues 389-397 from subunit A are well resolved and positioned in the gap between the AAA1 domains. **d)** Structural comparison of the linker domain conformations between the ATAD2B Walker B-ATPγS-H4K5acK8ac structure (yellow sticks) and the ATAD2-ATP structure (orange cartoon, PDB ID: 8h3h) to highlight the position of their linker domains (LD) between subunits F and A. The ATAD2 and ATAD2B structures were aligned by their AAA1 domains, and their F and A subunits are colored different shades of grey and blue. **e)** Structural comparison of the linker domain between the ATAD2B Walker B-ATPγS-H4K5acK8ac structure (yellow sticks) and the ATAD2-ATP-H3/H4K5Q structure, from state II in complex with the histone tetramer (red cartoon, PDB ID: 8juy). Alignment of the AAA1 domain highlights differences in the linker domain orientations. Density maps for the ATAD2B linker domain are shown in **Supplementary Figure 12**.

Structural comparison with the closely related human ATAD2 homolog shows that the ATAD2B LD adopts a trajectory similar to that observed in the ATP-bound ATAD2 complex, which also has an extended LD region wedged between the seam subunits (PDB ID: 8H3H, **Figure 7d**). However, the ordered LD in ATAD2B is seven residues shorter than the LD observed in ATAD2D, which reaches further into the seam nearly to the central pore^26^. To further contextualize their differences, we compare the ATAD2B LD with the LD conformations of ATAD2 bound to the histone H3/H4K5Q tetramer (PDB IDs: 8juw and 8juy). In these complexes, the LD is similarly truncated and displaced from the cleft (**Figure 7e**). Without the stabilizing interactions formed between the LD and the AAA1 domains of subunits A and F that lock ATAD2-ATP bound hexamer into a planar configuration, the histone bound state II displays greater flexibility.

When ATAD2 is bound to the histone H3/H4K5Q tetramer, a wider seam accompanies displacement of the LD, reflecting large rearrangements of the inter-subunit interface. Although ATAD2B was incubated with ATPγS and H4K5acK8ac prior to vitrification, the shorter histone peptide length and nucleotide state are not completely equivalent with the histone bound ATAD2 structures. Nevertheless, the partial ordering and seam localized positioning of the LD in ATAD2B resembles features of the histone-associated ATAD2 hexamers more closely than those of the ATP only state.

Collectively, these data demonstrate that ATAD2B contains and N-terminal linker domain that becomes ordered specifically at the hexameric seam, where it occupies the interface between the AAA1 domains of subunits A and F. The partial and asymmetric ordering of the LD segment, which is conserved in ATAD2, suggests the LD functions as a seam-specific stabilizing element.

### ATAD2B engages a peptide substrate through conserved pore loops

AAA+ ATPases translocate polypeptide substrates through a central pore formed by a ring of ATPase domains, using conserved pore loop elements arranged in a hexameric spiral^31^. The asymmetric AAA1 ring of ATAD2B shown in **Figures 3-5** suggests that its central pore may engage a substrate in a nucleotide state dependent manner. We examined the ATAD2B cryo-EM reconstruction for evidence of substrate binding and characterized how the conserved pore loop residues contribute to substrate engagement within the AAA1 domain.

During model building, we observed additional cryo-EM density within the central pore that does not correspond to any portion of the ATAD2B polypeptide chain (**Figure 8a and Supplementary Figure 13**). This density is confined to the AAA1 tier of the hexamer and is positioned within the central pore formed by the AAA1 ring. Similar density has been observed in the ATP-bound structures of ATAD2, Abo1, and Yta7 (PDB IDs: 8h3h, 6jpu,7uqj), where it has been hypothesized to correspond to a histone-derived substrate. In ATAD2B, nine residues were modeled into this density. Although the pore associated density is consistent with a polypeptide substrate and resembles substrate density observed in other ATAD2-like family members, insufficient side chain resolution precludes definitive assignment of substrate identity, and the density was therefore modeled as a polyalanine chain. (**Supplementary Figure 13a-b**). Coordination of the modeled substrate occurs through the pore loops of the AAA1 domains, which surround the polypeptide within the central pore (**Figure 8b,c and Supplementary Figures 1 and 13c**). Pore loop 1 (residues 474-481) is located between the β2 strand and α2 helix of the large nucleotide binding subdomain, while pore loop 2 (residues 511-519) lies between the β3 strand and α3 helix (**Figure 4a and Supplementary Figure 1**). In AAA+ ATPases pore loop 1 contains a conserved aromatic residue that mediates substrate interactions in ATP bound subunits^31^. In ATAD2B pore loop 1 residues Lys478-Trp479-Val480 wrap around the substrate backbone, and the conserved Trp479 residues from subunits B-E form a spiral staircase that tracks along the substrate within the pore (**Figure 8b,c**).

**Figure 8.**
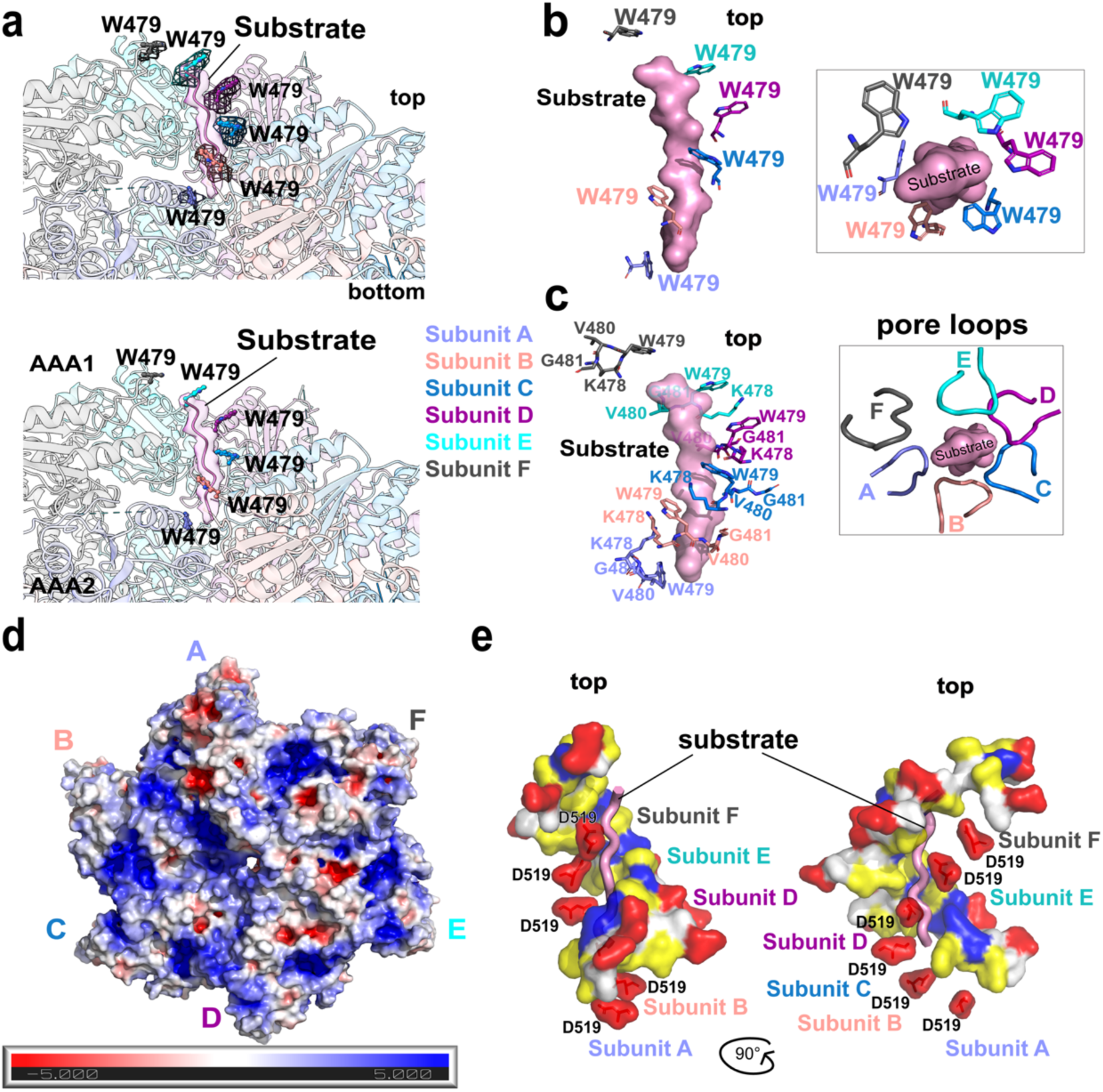
ATAD2B engages a polypeptide substrate within the central AAA1 pore. **a)** Top panel, cartoon representation of the ATAD2B Walker B hexamer colored by subunit, showing secondary structure elements. Cryo-EM density corresponding to a polypeptide substrate within the central pore of the AAA1 ring is shown in pink (contoured at 0.02 in ChimeraX), and the tryptophan staircase is shown as sticks, with the density for each residue (W479) in mesh. Bottom panel, cross section through the hexamer showing the substrate built into the density and positioned within the AAA1 pore. The substrate is shown as a cartoon, and the tryptophan staircase is shown as sticks. **b)** Side view (left) and top view (right) of the conserved tryptophan residues (W479, sticks) from each subunit forming a spiral staircase around the substrate (pink surface) within the central pore of the AAA1 ring. **c)** Side (left) and top (right) views of the AAA1 pore loop 1 residues (K478-G481, sticks) from each subunit arranged around the substrate, shown as a semi-transparent surface). **d)** Top view of the AAA1 central pore shown as a surface representation and colored by electrostatic potential (APBS calculation). **e)** Side views of the pore loop 1 residues (K478-G481) and pore loop 2 residue D519 from each subunit colored by residue type (yellow, hydrophobic; red, acidic; blue, basic; white, polar), illustrating the chemical environment of the pore surrounding the substrate (pink cartoon). Additional density maps for the tryptophan staircase and pore loops are found in **Supplementary Figure 13**.

In the ADP-bound subunit A at the base of the staircase, Trp479 is rotated downward and contacts the substrate at a greater axial separation, consisting with a disengaging state. In subunit F at the top of the spiral, Trp479 is positioned near the C-terminal of the peptide and poised to re-engage with the substrate as the cycle progresses (**Figure 8b,c**). As observed in other classical AAA+ ATPases such as Vps4, a lysine residue immediately preceding the conserved aromatic residue (Lys478) forms stabilizing cation-π stacking interactions with Trp479 within the same subunit and between neighboring subunits, reinforcing coordinated pore-loop motion (**Figure 8c)**^31,46^.

Pore loop 2 also adopts a spiral arrangement around the central pore. In subunit C, Asp519 forms a direct hydrogen bond with the substrate, while Asp519 from subunits D, E, and F are positioned close to the polypeptide, and those from subunits A and B lie below the N-terminal end of the modeled chain (**Figure 8e**). The axial distance between the Trp479 side chains in pore loop 1 and the Asp519 side chains in pore loop 2 that contact the substrate is approximately 7.5 Å, which corresponds to two residues along the polypeptide backbone. This spacing is consistent with a translocation step size of two amino acids, similar to other AAA+ ATPases including Yta7^25,47^.

In contrast to the substrate-engaged AAA1 ring, the corresponding pore loop regions in the AAA2 domains display distinct structural features and do not participate in direct substrate coordination. Notably, the AAA2 pore loops differ substantially from those of AAA1. AAA2 pore loop 1 (residues 806-809) lacks the conserved aromatic motif found in AAA1 and instead contains hydrophobic residues that point toward the central pore and are surrounded by hydrophilic residues acids that form an inter-subunit hydrogen bond network at the base of the pore. AAA2 pore loop 2 consists of a short Ala842-Val843-Ser844 motif connecting helices α3 and α4, which pack beneath the α1 and α2 helices connected by pore loop 1. The conserved Trp and Asp residues that directly contact the substrate in the AAA1 ring are absent in the AAA2 domain, and no cryo-EM density corresponding to substrate is observed within the AAA2 pore. Collectively, these features indicate that the AAA2 ring does not participate in substrate threading, but instead functions as a structural scaffold that stabilizes hexamer assembly and supports ATP-driven substrate translocation by the AAA1 domains.

Electrostatic surface an analysis of the ATAD2B hexamer reveals that the pore environment is optimized for substrate engagement **(Figure 8d**). A top-down view of the AAA1 domains shows that the surface displays a heterogenous charge distribution across the hexameric ring, with distinct acid and basic patches arranged in a subunit specific pattern. The entrance to the central pore is lined by acidic and basic electrostatic potential, with more negative patches inside the central pore. Side view representations indicate that this electrostatic potential arises from residues flanking the pore loops including Asp519 of pore loop 2, while residues such as Trp479 and Lys478 make direct hydrophobic contacts with the substrate. Together the mixed electrostatic surface of AAA1 and the more negative chemical environment of the pore suggests that substrate entry may be guided toward the pore by electrostatic complementary, and ATP driven pore loop motions provide the direct mechanical interactions required for processive substrate translocation.

Together, these data demonstrate that ATAD2B engages a polypeptide substrate specifically within the central pore of the AAA1 ring through conserved pore loop elements arranged in an asymmetric spiral staircase. Substrate coordination is mediated by pore loops 1 and 2 from ATP-bound AAA1 subunits, whereas the corresponding pore loop regions of the AAA2 domains lack substrate interacting features and do not participate in substrate threading, consistent with their non-catalytic role. The defined geometry and spacing of pore loop contacts support stepwise substrate translocation through the AAA1 pore during the ATPase cycle.

These structural observations establish a mechanistic framework for substrate engagement by ATAD2B and provide a foundation for understanding how asymmetric ATP hydrolysis, seam organization, and multiple stabilizing elements are coordinated to drive substrate processing.

## Discussion

In this study, we establish human ATAD2B as a mechanochemically active member of the classical type II AAA+ ATPase family and define the structural principles underlying its ATP-driven function. The cryo-EM structure reveals that ATAD2B assembles into a stable two-tiered hexamer in which an asymmetric AAA1 spiral staircase engages and coordinates substrate within the central pore, while a planar, catalytically inactive AAA2 ring provides structural stabilization. This architecture is accompanied by asymmetric nucleotide occupancy, with flexibility localized at the seam, and conserved inter-subunit signaling features that together support a model of sequential ATP hydrolysis coupled to stepwise substrate translocation. In contrast to its paralog ATAD2, which exhibits minimal ATPase activity, ATAD2B displays robust ATP hydrolysis and conserved pore loop engagement, placing it mechanistically closer to the ATAD2-like yeast homologs Abo1 and Yta7. These findings provide a structural framework for understanding how ATAD2B couples ATP hydrolysis to substrate processing and establish a foundation for further investigation into its distinct chromatin associated functions.

### ATP-driven substrate translocation by an asymmetric AAA1 ring

The shallow spiral staircase formed by the AAA1 domains of ATAD2B provides a structural framework for understanding how ATP hydrolysis is coupled to substrate engagement and translocation (**Figures 3 and 8**). In AAA+ ATPases, the asymmetric ring architectures is a hallmark of motors that hydrolyze ATP sequentially around the hexamer, producing directional movement of substrates through a central pore^31,35^. The AAA1 ring of ATAD2B adopts this conserved organization, with a cumulative vertical offset across subunits and increased flexibility localized to subunits A and F at the seam interface (**Figures 3a-c**). This architecture, together with the asymmetric nucleotide occupancy observed across the ring (**Figure 4**), supports a rotary ATPase mechanism in which individual subunits cycle through ATP-bound, hydrolyzing, and ADP-bound states rather than acting synchronously.

Direct evidence for substrate engagement by ATAD2B is provided by the presence of additional density within the central pore of the AAA1 ring (**Figure 8a-c**), which does not correspond to any region of the ATAD2B polypeptide chain and is therefore consistent with a polypeptide substrate. Similar pore associated density has been reported in ATP-bound structures of ATAD2-like family members Abo1, Yta7, and ATAD2, where it has been attributed to histone-derived substrates or histone tails^22,25,26^. In ATAD2B, this density is confined to the AAA1 tier of the hexamer and is coordinated by conserved pore-loop elements, indicating that substrate contact and mechanical work are restricted to the catalytically active AAA1 ring.

Substrate coordination within the AAA1 pore is mediated by pore loop 1 and pore loop 2 residues that form a spiral staircase aligned with the AAA1 asymmetry (**Figure 8c,e**). The conserved aromatic residue Trp479 of pore loop 1 intercalates along the substrate backbone in ATP-bound subunits, forming a continuous path of engagement through the pore. This arrangement mirrors the tryptophan staircases observed in other type II classical AAA+ ATPases, including NSF and Pex1/Pex6, which are known to translocate substrates through processive, hand-over-hand mechanisms^34,48,49^. In ATAD2B, the conformation of Trp479 differs across subunits. ATP-bound subunits with interlocking interfaces form the principal substrate gripping interfaces, whereas the ADP-bound seam subunit A adopts a disengaging configuration, with Trp479 rotated away from tight substrate contact (**Figure 8b,c**). The other seam subunit F, located at the top of the staircase, positions Trp479 near the substrate at the entrance to the pore, consistent with a poised re-engagement state as the catalytic cycle progresses.

Pore loop 2 further reinforces substrate coordination through a negatively charged aspartate residue that likely optimizes the interaction with positively charged histone proteins^50^. Asp519 from pore loop 2 forms direct contacts with the substrate in subunits located in the middle of the staircase (C/D) (**Figure 8e)**. Equivalent residues in subunits lower in the spiral (A/B) are positioned below the N-terminal end of the modeled polypeptide, while the sidechains of Asp519 in subunits E-F are reaching upwards toward the substrate. The residues in pore loop 2 are not as well conserved among AAA+ ATPases, as they are in pore loop 1, but they appear to contribute to substrate specificity^31^. For example, in the Spastin AAA+ ATPase mutations in a basic residue of pore loop 2 abrogates binding with negatively charged tubulin, and an aromatic residue in pore loop 2 of YME1 contributes to the degradation rate of hydrophobic membrane substrates of this AAA+ protease^51,52^. All members of the ATAD2-like AAA+ ATPase family contain a negatively charged residue in pore loop 2 (**Supplementary Figure 1**), and in Yta7, Glu532 of subunits E and F makes a direct hydrogen bond contacts to lysine residues in the histone H3 substrate found in the pore^25^. Thus, the negatively charged pore loop 2 appears to play an important role in stabilizing histone substrates in the central pore, which are enriched in positively charged lysine and arginine residues^53^.

Importantly, substrate is not observed within the AAA2 ring of ATAD2B, and the corresponding AAА2 pore loop regions lack the conserved aromatic and acidic residues that mediate substrate interactions in the AAA1 ring (**Figure 8a-c and Supplementary Figure 13**). Instead, the AAA2 ring remains planar and highly symmetric (**Figure 3**), indicating that this tier does not directly participate in substrate threading or mechanical translocation. This division of labor between the AAA1 and AAA2 domains represents a defining organizational feature of ATAD2B and distinguishes it from many other type II AAA+ ATPases that contain classical AAA+ domains in tandem^21^. For example, in NSF and p97, both AAA+ ATPase rings bind nucleotides and contribute to substrate contacts in the central pore^41,48^. Only a limited number of AAA+ systems share an architecture consisting of tandem AAA+ domains with a catalytically inactive module, notably the Pex1/Pex6 complex. However, in that system the first AAA+ domain (D1) in Pex1 and Pex6 are inactive and serve a primarily structural role, and these two proteins function together in a hetero-hexameric assembly^36,54^. In ATAD2B, confinement of catalytic activity to the AAA1 ring supports a model in which AAA2 functions as a non-catalytic structural scaffold that stabilizes the global hexameric assembly during asymmetric ATP hydrolysis and substrate translocation.

Electrostatic surface analysis further supports a specialized role for the AAA1 pore in substrate engagement. The surface of the AAA1 domains exhibits a heterogeneous charge distribution, with acidic residues lining the pore entrance and inner walls, creating a negatively biased electrostatic environment that would favor accommodation of basic polypeptide substrates such as histone tails (**Figure 8d**). These long-range electrostatic features likely guide histone substrates toward the pore, while direct hydrophobic and polar contacts mediated by pore loop residues provide the ATP dependent grip required for mechanical translocation. Together, the asymmetric organization of the AAA1 ring, conserved pore-loop architecture, and electrostatically tuned pore environment place ATAD2B squarely within the mechanistic framework of classical AAA+ protein remodeling complexes^21,31,55^.

Collectively, these observations support a model in which sequential ATP hydrolysis propagates conformational changes around the AAA1 ring, coupling nucleotide state to pore loop geometry and enabling stepwise substrate movement through the central channel (**Figures 3-5 and 8**). While ATAD2B contains several unique stabilizing features not found in other AAA+ motors, its core mechanism of ATP-driven substrate translocation appears to be evolutionarily conserved, providing a basis for understanding how this previously uncharacterized human AAA+ ATPase contributes to chromatin associated processes. It should be noted that the structure was determined in the presence of a short, acetylated histone H4 peptide rather than a full histone complex or nucleosome. While this ligand likely approximates a physiologically relevant substrate, interactions with intact chromatin would impose additional steric or regulatory constraints not captured in the current structure.

### Comparisons within the ATAD2-like family

Members of the ATAD2-like protein family, which includes ATAD2B, ATAD2, Yta7, and Abo1, all share a conserved mechanochemical core and display unique architectural adaptations that distinguish them from other type II classical AAA+ ATPases. All four proteins assemble into a two-tiered hexameric complex formed by their tandem AAA+ domains, with ATP hydrolysis confined to their asymmetric AAA1 ring, and a planar, catalytically inactive AAA2 ring provides structural support^22,25,26^. This conserved organizational principle underlies their ability to engage histone substrates through pore loops in a nucleotide dependent fashion. However, unlike most type II AAA+ ATPases, the ATAD2-like proteins do not appear to rely on nucleotide binding in both ATPase tiers for complex stability, suggesting that alternative stabilizing mechanisms have evolved to support hexamer formation and catalytic activity.

One striking and unique feature shared by all ATAD2-like family members is the presence of a noncanonical stabilizing insertion within the AAA2 domain that forms the knob region, which is followed by an unstructured loop that connects AAA2 to the bromodomain (**Figure 6 and Supplementary Figure 11**). In Abo1 and Yta7, these insertions are well ordered and form extensive inter-subunit contacts that stabilize the hexamer even under nucleotide-free conditions^22,25^. The knob-hole architecture has not been observed in other AAA+ ATPase families, and appears to serve as an alternative means of stabilizing the hexameric assembly in lieu of nucleotide dependent stabilization of the AAA2 ring as observed in NSF, p97, ClpB, and Hsp104^27,33,48,56^. Sequence and structural analyses indicate that analogous insertions are present in ATAD2 and ATAD2B, although in this context, ATAD2B occupies an intermediate position between its paralog ATAD2 and the yeast homologs Abo1 and Yta7. The Cryo-EM reconstruction of ATAD2B shows slightly less well resolved knob-hole and linker arm density, while ATAD2 exhibits weaker ordering and reduced hexamer stability, suggesting that the stability conferred by these features is incomplete or more tightly regulated (**Figure 6 and Supplementary Figures 1 and 11**)^26^. These observations suggest that the knob-hole and linker arm features represent a family-specific adaptation that compensates for the lack of ATP binding in AAA2, enabling fine-tuning of oligomer stability without relying on canonical nucleotide dependent mechanisms.

### Functional divergence across the ATAD2-like family

Despite sharing a conserved AAA+ mechanochemical core, members of the ATAD2-like family perform distinct chromatin-associated functions that are reflected in differences in their ATPase activity, substrate engagement, and chromatin targeting. Abo1, Yta7, ATAD2, and ATAD2B all associate with histone substrates as evidenced by biochemical and structural studies spanning the yeast and human homologs^2,6,22,57,58^. However, how ATP hydrolysis is coupled to histone handling differs markedly among family members. For example, Abo1 facilitates deposition of H3-H4 histones onto DNA, and promotes recycling of FACT, consistent with a role in nucleosome assembly^22,23,59^. In contrast, Yta7 evicts histone H3 from nucleosomes, contributes to nucleosome remodeling during DNA replication, and also helps Scm3 load the centromeric histone H3 variant Cse4 in yeast^24,25,60^. The chromatin associated roles of ATAD2 and ATAD2B are less well defined. ATAD2 interacts with chromatin where it functions as a transcriptional co-activator by regulating the local availability of the histone chaperone HIRA, which deposits the histone H3.3 variant in chromatin^6,61,62^. ATAD2 and ATAD2B bind to newly synthesized H4 histones that are di-acetylated at H4K5acK12ac, which may serve to protect against histone deacetylase activity until they are deposited into chromatin, but how ATAD2 and ATAD2B contribute to nucleosome assembly and/or disassembly is currently unknown^2,5,63^.

An additional layer of functional divergence within the ATAD2-like family arises from differences in their bromodomains and N-terminal domains (NTDs). Abo1 and Yta7 harbor non-canonical bromodomains that preferentially bind unmodified histone H3, while ATAD2 and ATAD2B contain canonical bromodomains that recognize specific combinations of acetylation modifications on the H4 and H2A histones^3,22,25^. The ATAD2B bromodomain binds to a broader range and combinations of acetyllysine modifications than ATAD2, including those on the histone H2A variants, H2A.X and H2A.Z, which may reflect cell cycle specific roles for these human homologs^23^.

Another striking difference among the ATAD2-like family arises from the organization and function of their NTDs. In many AAA+ ATPases the N-terminal regions play critical roles in substrate recognition, mediate protein-protein interactions with chaperones or accessory proteins, and can tether them to membranes^31^. Within the ATAD2-like family, the N-terminal domains are poorly conserved at the sequence level, but they appear to play key roles in chromatin association and functional specificity. In the yeast homologs, cryo-EM reconstructions of Yta7 show that it contains a structured N-terminal domain that cooperates with the bromodomain to form a third tier above the AAA+ rings, generating a distinct histone-binding platform essential for nucleosome recognition and remodeling^25^. This N-terminal architecture is functionally integral to Yta7’s ability to evict histone H3 from nucleosomes and to remodel chromatin during replication and centromere maintenance^57,60^. In Abo1 N-terminus also appears to be functionally important as deletion of the NTD abolishes its interaction with the FACT complex and compromises its histone-loading activity, despite leaving ATP hydrolysis largely intact^22,59^.

These findings suggest that, in yeast homologs, the NTD act as critical interfaces for mediating interactions with other proteins, rather than as components of the ATPase motor itself.

The NTD of human ATAD2 contains a D/E repeat enriched in aspartic and glutamic acid residues that enhance its interaction with positively charged histones, and is required for the association of ATAD2 with chromatin^64^. This is consistent with data on the original short form of ATAD2 (ΔN-ATAD2) found in testis, which has a decreased ability to bind to chromatin^32^.

Previous studies on ATAD2 also found that the NTD is required for its co-activator functions by mediating interactions with the androgen receptor, MYC, and E2F^7,8,65^. ATAD2B remains less well characterized with respect to its N-terminal domain. Similar to ATAD2, the N-terminal region of ATAD2B is predicted to be largely unstructured and it contains a region enriched in acidic residues. However, the N-terminal domain of ATAD2B is 19 residues shorter than in ATAD2 (residues 1-412 versus 1-431, respectively) and the acidic region of ATAD2B is smaller and enriched in glutamate residues, when compared to the long stretch of aspartates found in the ATAD2 N-terminus (**Supplementary Figure 1**). The ATP hydrolysis rate of WT ΔN-ATAD2B a 0.34 ATP/hexamer/sec is substantially higher than that of ATAD2, which was nearly inactive, yet modestly reduced relative to the yeast homologs (**Figure 1c and Supplementary Figure 2c**)^22,25,26^. Unlike ATAD2, truncation of the N-terminal domain in ATAD2B does not abolish ATP hydrolysis. This suggests that the N-terminus of ATAD2B may play a more modulatory or context-dependent role, potentially contributing to chromatin targeting or regulatory interactions, without being strictly required for motor activity. At present however, the functional contribution of the ATAD2B N-terminus remains unresolved.

Importantly, a limitation of the present structural analysis of ATAD2B, and prior cryo-EM studies on ATAD2 and Abo1, is the use of N-terminally truncated constructs to facilitate biochemical stability of the hexameric complex for structure determination^22,26^. As a result, direct observation of the N-terminal domain and its association with the AAA+ core is not possible in these structure. Given the established role of the N-terminal regions of Yta7, Abo1, and ATAD2 in chromatin engagement, the absence of this domain in the current ATAD2B structure precludes definitive conclusions regarding how ATP hydrolysis and pore loop substrate engagement are coupled to chromatin binding *in vivo*. Determination of full-length structures of ATAD2B, particularly in complex with intact chromatin substrates, will be essential to resolve how its N-terminal domain contributes to function.

In conclusion, this study presents the first high-resolution cryo-EM structure of human ATAD2B and establish it as a mechanochemically active member of the ATAD2-like AAA+ ATPase family. The structure reveals a conserved yet tunable ATPase architecture in which an asymmetric AAA1 ring hydrolyzes ATP sequentially and coordinates substrate engagement through conserved pore-loop elements, while a planar, non-catalytic AAA2 ring provides a stable architectural scaffold. Together with biochemical measurements of ATP hydrolysis, these findings demonstrate that ATAD2B is not a passive paralog of ATAD2 but instead functions as an ATP-dependent molecular machine capable of engaging chromatin substrates.

A comparative analysis indicates that functional divergence among ATAD2-like proteins arises primarily from differences in ATP hydrolysis rates, and extent of substrate engagement within the AAA1 pore, while auxiliary domains peripheral to the AAA+ core, such as the bromodomain and N-terminal domain, are likely involved in regulation and chromatin targeting specificity. Yeast homologs Abo1 and Yta7 apply the conserved AAA+ motor to histone loading and nucleosome remodeling, respectively. While the chromatin associated functions of human ATAD2 and ATAD2B are currently unknown, the combination of a canonical bromodomain and an active AAA1 motor within a stable hexameric ATAD2 assembly, is consistent with an active role in ATP-dependent chromatin associated activities ranging from remodeling to regulated histone chaperoning. These results provide a mechanistic framework for the rational design of therapeutic strategies targeting this clinically relevant chromatin regulator in cancer, respiratory and neurological diseases^1,12^.

Looking forward, determination of full-length ATAD2B structures in complex with intact histone assemblies or nucleosomes will be critical for understanding how ATP-driven substrate threading is coupled to the functions of the N-terminal region and bromodomain guided histone recognition. Mutational analysis of stabilizing insertions, pore-loop residues, inter-subunit signaling motifs, and the bromodomain binding pocket, combined with cellular and genomic assays, will further clarify how ATAD2B contributes to transcriptional regulation, genome maintenance, and DNA repair. More broadly, the architecture revealed here provides a molecular basis for understanding how the ATAD2-like family repurposes a conserved AAA+ motor through unique stabilization strategies and modular chromatin-binding elements to expand the functional diversity of AAA+ ATPases in chromatin biology.

## Methods

### Plasmid construction and protein overexpression

Human wild-type (WT) ATAD2B (residues 380-1458, UniProt ID: Q9ULI0) and human ATAD2B with a Walker B mutation (E506Q) were cloned into a pFastBac1 plasmid containing an N-terminal GST-tag, followed by a PreScission Protease Cleavage site and a His6x-tag followed by the ATAD2B sequence which was codon optimized and synthesized by GenScript (Piscataway, NJ, USA). Each pFastBac1 plasmid was transformed into DH10Bac competent cells (gift from Dr. Kathleen Trybus). Recombinant bacmids were selected by blue-white screening as indicated in the Bac-to-Bac baculovirus expression system (Thermo Fisher Scientific, Waltham, MA, USA)^66^. Each bacmid was used to infect Sf9 cells, and each protein required specific optimization of amplification. For the human ATAD2B WT, the virus was amplified three times, for a final multiplicity of infection of (MOI) of 23. For the human ATAD2B Walker B mutant, the virus was amplified two times, for a final MOI of 15. For each protein, 3 billion cells were infected for 2 hours, and replaced with media to grow for 72 hours (ATAD2B Walker B) or 96 hours (WT ATAD2B).

### Protein Purification

After the infection period, the cells were harvested by centrifugation at 5,000 RPM for 10 minutes at 4°C. The cell pellets were resuspended in 45-60 mL of lysis buffer (WT ATAD2B: 50 mM Tris, pH 7.5, 150 mM NaCl, 5% glycerol, 10 mM MgCl_2_, 1 mM dithiothreitol (DTT), 2 mM ATP; ATAD2B Walker B: 50 mM Tris, pH 7.5, 150 mM NaCl, 5% glycerol, 2 mM MgCl_2_, 1 mM dithiothreitol (DTT), and the lysis buffer was supplemented with 100 mM PMSF and 1 protease inhibitor tablet (Pierce protease inhibitor tablets, EDTA-free, Thermo Fisher Scientific). The cells were lysed by sonication, which was clarified by ultracentrifugation at 50,000 RPM for 30 minutes at 4°C. The supernatants were incubated with 1.5-2 mL of glutathione agarose resin (Thermo Fisher Scientific,) for 1 hour, 45 minutes with gentle agitation at 4°C. The suspension was poured into 10-mL Polypropylene columns (Thermo Fisher Scientific) and washed with 2 CV of Wash Buffer (ATAD2B WT: 50 mM Tris, pH 7.5, 150 mM NaCl, 5% glycerol, 10 mM MgCl_2_, 1 mM DTT, 2 mM ATP; ATAD2B Walker B: 50 mM Tris, pH 7.5, 150 mM NaCl, 5% glycerol, 2 mM MgCl_2_, 1 mM DTT). The GST-tagged WT and Walker B ATAD2B proteins were eluted off the beads with 20 mM reduced glutathione. The WT ATAD2B was dialyzed into Wash Buffer and ATAD2B Walker B was step-dialyzed into cryo-EM buffer, changed after three hours (20 mM HEPES pH 7.5, 150 mM NaCl, 5% glycerol, 2 mM MgCl_2_, 1 mM DTT) to remove the reduced glutathione overnight. Post-dialysis, the proteins were concentrated: WT ATAD2B: 7.87 mg/mL and ATAD2B Walker B: 12.48 mg/mL, using an AmiconR Ultra (MILLIPORE) with a 10-30 kDa MW cut-off for size exclusion chromatography (SEC) using a Superdex 200 Increase 10/300 GL column on an ÄktaPrime system (GE/Cytiva) in Wash Buffer (WT ATAD2B) or cryo-EM buffer (ATAD2B Walker B). Peaks corresponding to each ATAD2B construct were pooled and concentrated to a final concentration of WT ATAD2B: 2.3-2.5 mg/mL (15.2-16.8 µM), ATAD2B Walker B: 4.87 mg/mL (32.4 µM) and flash frozen in liquid nitrogen and stored at-80 °C for future experiments. Protein purity was confirmed by sodium dodecyl sulfate-polyacrylamide gel electrophoresis (SDS-PAGE) gels stained with Bio-Safe Coomassie G-250 Stain (Bio-Rad). Protein concentrations were determined with NanoDrop (Thermo Fisher Scientific).

### Histone peptide synthesis

The histone H4K5acK12ac (residues 1-24) peptide was purchased from GenScript (Piscataway, NJ, USA). The amino acid sequence is: SGRG(Kac)GGKGLG(Kac)GGAKRHRKVLRD. This peptide was synthesized with N-terminal acetylation (N*α*-ac) and C-terminal amidation and was >95% pure confirmed by high-performance liquid chromatography analysis. Mass Spectrometry was used to confirm their identity.

### Mass Spectrometry

Protein identification was carried out using liquid chromatography / mass spectrometry (LC/MS). Purified ATAD2B fractions were electrophoresed on SDS-PAGE, and individual Coomassie Blue-stained bands of interest were excised, minced to approximately 1 mm^3^ cubes, followed by the standard in-gel digestion protocol. The dried digests were resuspended in 2.5% acetonitrile (ACN) / 0.1% formic acid (FA) and analyzed by nanoflow ultra-performance LC/MS on the EASY nLC 1200 coupled to an Orbitrap Exploris 240 mass spectrometer (Thermo Fisher Scientific). Samples were loaded onto a 100 μm x 300 mm capillary fused silica column that was constructed by laser pulling its end to a ∼3 μm orifice, making an integral frit by packing with minimal amount of 5-μm Magic C18AQ, and filling the column with 1.8-µm 120 Å Uchrom C18 chromatography resin (nanoLCMS Solutions, CA, USA). Peptides were separated by a solvent system composed of solvent A: 100% water / 0.1% FA and solvent B: 80% ACN / 0.1% FA at a flow rate of 300 nL min^-1^ with a gradient of 2-25% B over 30 min, 25-40% B in 10 min, 40-95% B in 5 min, then 95% B for 4 min, followed by an immediate return to 50% B in 1 min and a hold at 5% B over 5 min. Peptides were introduced into the mass spectrometer via the Nanospray Flex ionization source with a spray voltage of 1.9 kV, an ion transfer tube temperature of 300 °C, and RF lens set at 70%. “Top speed in 2 seconds” acquisition mode was used with peak width set as 20 s for automatic gain control (AGC) estimation. Survey scans were collected for *m/z* 350 - 1400 at 60,000 resolution (AGC target: 250%; max IT 25 ms; profile mode) and data-dependent higher energy collisional dissociation tandem mass spectrometry scans (AGC target: 50%; Max IT: Auto; centroid mode) were acquired at 15,000 resolution with quadrupole isolation at an isolation width of 2 *m/z* and a normalized collision energy at 30%. Intensity threshold was set as 5e^3^, and dynamic exclusion was enabled (exclusion duration: 30 s). Mass spectrometry data were analyzed using Proteome Discoverer 2.5 (Thermo Fisher Scientific). Product ion spectra were correlated to known sequences by SEQUEST in the processing workflow against the *Spodoptera frugiperda* (UP000829999; downloaded on Mar. 6, 2025) and *Homo Sapiens* (UP000005640; downloaded on Nov. 8, 2024) UniProt fasta databases. Search parameters were as follows: (1) full trypsin enzymatic activity; (2) maximum missed cleavages = 2; (3) minimum peptide length = 6; (4) mass tolerances set at 10 ppm for precursor ions and 0.02 Da for fragment ions, respectively; (5) dynamic modifications on methionines (+15.9949 Da: oxidation) and protein termini (+42.01 Da: acetyl; - 131.040 Da: met-loss;-89.030 Da: met-loss+acetyl); (6) 4 maximum dynamic modifications were allowed per peptide; and (7) static carbamidomethylation modification on cysteines (+57.021 Da). Percolator node was included in the workflow to limit the false discovery rate to less than 1% in the datasets with the use of a concatenated database.

### Negative Stain Electron Microscopy (EM)

Continuous carbon-coated mesh copper grids (Ted Pella, Inc.) were glow discharged with a Gatan Solarus Model 950 Advanced Plasma System for 50 s with H_2_ and O_2_ gas. The SEC-purified WT ATAD2B and ATAD2B Walker B proteins were diluted to ∼0.1 mg/mL and ∼0.05 mg/mL, respectively. Two samples were created for each, one with ATP added to a final concentration of 2 mM, and one without ATP. Then 3.0 µL of each protein sample was added to the glow discharged grid and incubated on the grid for 30 seconds. Excess protein was blotted off by touching the grid edge to filter paper. WT ATAD2B grids were washed three times with 20 µL drops of Milli-Q H_2_O, and ATAD2B Walker B grids were washed two times with 20 µL drops of Milli-Q H_2_O. The grids were then stained by dipping the grids sequentially into three 20 µL drops of 2-3% uranyl acetate, and incubating for 30 seconds in the last drop. The excess stain was removed by blotting with filter paper. The grids were allowed to dry on the bench and stored in a desiccator overnight until imaging. The grids were observed on a 120kV JEOL 1400 transmission electron microscope (TEM) at 20,000-60,000 x located in the Microscopy Imaging Core (MIC) at the University of Vermont. Micrographs of the ATAD2B WT and Walker B samples, with and without ATP, were collected with a CCD camera (AMT XR611 high resolution 11 megapixel).

### ATPase Assays

The EnzChek Phosphate assay kit (Thermo Fisher Scientific) was used to measure the ATPase rate of ATAD2B. Briefly, SEC-purified and concentrated WT and Walker B ATAD2B was thawed and diluted to a final concentration of 0.25 µM and incubated with the kit’s reagents in our ATPase Assay Buffer (50 mM Tris pH 7.5, 150 mM NaCl, 5% glycerol, 10 mM MgCl_2_, and 1 mM DTT) in a 1.0 mL volume at room temperature for 10 minutes prior to the addition of ATP. ATP was added to the reaction in a 96-well clear bottomed plate (Corning) to a final concentration of 2 mM and absorbance was immediately read at 360 nm on a SpectraMax M4-R (Molecular Devices) plate reader at 37°C. Thereafter, absorbance measurements were taken every 10 seconds for 15 minutes, and steady state was defined when the rate was linear, approximately from 210-690 seconds. The enzymatic rate was calculated from each experiment, which were performed at least in triplicate to obtain an average ATP hydrolysis rate, which is reported with the standard deviation value.

### Cryo-EM grid preparation and data collection

UltrAuFoil grids 300 mesh R1.2/1.3 were glow discharged using a Gatan Solarus II with the grid bar side facing up with Ar_2_ and O_2_ for 10 seconds, twice. The SEC-purified and concentrated ATAD2B Walker B protein samples were thawed and prepared for vitrification.

The ATAD2B Walker B protein (final concentration: 4.09 mg/mL, 27.2 µM) in cryo-EM buffer was incubated with 2 mM adenosine 5’-(-thio)-triphosphate tetra-lithium salt (ATPγS, Cayman Chemical), histone peptide H4K5acK12ac (residues 1-24) in a 1:10 ratio (272 µM), and 0.025% beta-octyl-glucoside in a 10 µL total volume. Afterwards, the complex was incubated for ∼30 minutes on ice prior to vitrification. For freezing, 3 µL of this ATAD2B-ATPγS-H4K5acK12ac protein complex was frozen on glow discharged grids using a Vitrobot Mark IV (Thermo Fisher Scientific) set to 4 °C with 0 blot force, and a blotting time of 1 second. These grids were frozen at the National Center for Cryo-EM Access and Training, and shipped to the National Cancer Institute’s National Cryo-EM Facility at the Frederick National Laboratory for Cancer Research where 4,870 movies were collected on a Titan Krios 300 kV microscope with a Gatan K3 detector and defocus range of 0.8-2.5 µm, with a 0.2 µm step (**Supplementary Table 1**).

### Cryo-EM Data Processing

All data processing was performed in cryoSPARC v.4.5.3^67^. As shown in **Supplementary Figure 3**, 4,870 movies were pre-processed through motion correction with patch motion correction. Patch contrast transfer function (CTF) estimation was done on the motion-corrected micrographs. These pre-processed micrographs (**Supplementary Figure 4a)** were then manually screened, and 201 micrographs were discarded due to poor ice quality, bad CTF estimations, or the resolution cutoff of 9 Å. A previous dataset was used to generate 10 templates (three top/bottom, four side, two tilted) that were subsequently used for template picking **(Supplementary Figure 4b)**. Template picker picked 2,669,568 particles from the remaining 4,573 micrographs. We used the inspect particle picks job to filter through these picks based on defocus-adjusted power and pick scores, ending with 1,484,271 particles. The particles were extracted with a box size of 484 pixels and binned by 2. The resulting 1,363,273 particles were subjected to two rounds of reference-free two-dimensional (2D) classification, where 779,108 protein particles were separated from junk. The protein class and junk class were each used to make 3D volumes for filtering in 3D with three successive rounds of heterogeneous refinement, after which 215,307 particles were left. These particles were re-extracted at the full box size of 484 pixels, and the remaining 213,605 particles (without binning) were used in an initial round of homogenous refinement. Next, particle subtraction was carried out and an inverted mask was built around the AAA+ ATPase domains to exclude the flexible, disordered bromodomain to ultimately improve alignments. Two rounds of Global CTF refinement were carried out, followed by non-uniform refinement without a mask. Finally, reference-based motion correction on the curated set of 213,605 particles, followed by non-uniform refinement was used to produce the final map with a resolution of 3.0 Å according to the 0.143 gold standard FSC cutoff criteria (**Supplementary Figure 4c,d)**.

### Atomic model building and refinement

The AlphaFold model of monomeric ATAD2B (AF-Q9ULI0-F1-v4) was downloaded using the Process Predicted Model module in Phenix, where its low-confidence predicted loop regions were trimmed and bromodomain was manually removed from the.pdb file^44,68^. Six copies of this processed AlphaFold ATAD2B model were docked into the cryo-EM map by individually aligning each model to a chain in the previously solved ATAD2 structure (PDB ID: 8h3h)^26^. The was model fit into the map using rigid body refinement, simulated annealing, and morphing in Phenix. Multiple rounds of refinement in Phenix were used in addition to manual model building in Coot to prepare the final structure^69–72^. During the refinement, secondary restraints were applied. Correlation coefficients (model-to-map fit in Phenix) were closely monitored to avoid overfitting of the model in the map.

### Accession numbers

Will be made available after acceptance.

## Acknowledgments

We gratefully acknowledge the staff at the National Center for Cryo-EM Access and Training (NCCAT) at the New York Structural Biology Center for their exceptional support and guidance on sample preparation, data collection, and analysis for structure determination of ATAD2B using single-particle cryo-electron microscopy. We thank Wai Lam for analysis of the mass spectrometry data. We are especially grateful to Dr. Kathleen Trybus at UVM for assistance with protein expression using the Sf9 insect cells. We thank Dr. Zhiqing (Jenny) Wang at the National Cryo-EM Facility at the Frederick National Laboratory for Cancer Research for data collection on the ATAD2B-ATPγS-H4K5acK12ac complex. We thank members of the Glass laboratory for critically reading the manuscript.

## Funding information

This work was supported by a mid-career advancement award from the Molecular and Cellular Biosciences Division of the National Science Foundation to KCG and MAC under award number 2321501. This work was also supported by the National Cancer Institute and the National Institute of General Medical Sciences of the National Institutes of Health under award numbers P01CA240685 and R01GM129338 to KCG, and R35GM136288 to KT. The Vermont Advanced Computing Center (VACC) is supported by the National Science Foundation under award number 1827314. Creation of the UVM Center for Biomedical Shared Resources, which houses the Microscopy Imaging Center, was supported by NIH award 1C06OD030087-01. The Vermont Biomedical Research Network Proteomics Facility (RRID: SCR_018667) is supported through NIH grant P20GM103449 from the INBRE Program of the National Institute of General Medical Sciences. Any opinions, findings, and conclusions or recommendations expressed in this material are those of the author(s) and do not necessarily reflect the views of the National Institutes of Health or the National Science Foundation. This research was also supported by the University of Vermont Cancer Center, and the University of Vermont Larner College of Medicine. Some of this work was performed at the National Center for Cryo-EM Access and Training (NCCAT) and the Simons Electron Microscopy Center located at the New York Structural Biology Center, supported by NIH (Common Fund U24GM129539, NIGMS R24GM154192), the Simons Foundation (SF349247), and the NY State Assembly. Part of the cryo-EM work was also supported by the National Cancer Institute’s National Cryo-EM Facility at the Frederick National Laboratory for Cancer Research under Contract 75N91019D00024.

## Author Contributions

KLM, MAC, KMT, and KCG conceptualized the work. KLM, JML, PMF, JEM, and KCG performed research. KLM, EYDC, JML, PMF, JEM, KMT, MAC, and KCG analyzed data. KLM and KCG wrote the paper, and all authors revised and approved the manuscript. All authors approved the submission and agreed to be responsible for their contributions.

## Competing Interests

The authors declare no competing interests.

## Supplemental Information

**Supplementary Table 1.** Cryo-EM data collection, refinement, & validation statistics.

| Data Collection and Processing |  |
| --- | --- |
| EMDB Code | *available after acceptance* |
| PDB Code | *available after acceptance* |
| Electron Microscope | Titan Krios |
| Camera | Gatan K3 |
| Magnification | 105000x |
| Voltage | 300 kV |
| Nominal defocus range ($\mu\text{m}$) | -0.8 to -2.5 |
| Total electron dose ($\text{e}/\text{\AA}^2$) | 50.03 |
| Pixel Size ($\text{\AA}$) | 0.835 |
| Exposure time (sec) | 23.3 |
| Number of frames per movie | 50 |
| Automation software | Latitude S ver. 3.44 (Gatan) |
| Micrographs collected | 4,870 |
| Data processing software | cryoSPARC v.4.5.3 |
| Symmetry imposed | None: Default C1 |
| Total extracted particles | 1,363,273 |
| Total refined particles | 213,605 |
| Final particles | 213,605 |
| Map resolution (FSC=0.143/$\text{\AA}$) | 3.0 |
| Local resolution range ($\text{\AA}$) | 3-6.5 |
| Refinement |  |
| Initial Model (AlphaFold code) | AF- Q9ULI0-F1-v4 |
| Refinement package | Phenix 1.21.2-5419 |
| Model resolution (FSC=0.5/$\text{\AA}$) | 3.2 |
| Map sharpening B factor ($\text{\AA}^2$) | NA |
| Map CC | 0.86 |
| Model composition |  |
| Non-hydrogen atoms | 26769 |
| Protein residues | 3301 |
| Ligands | 6 |
| B factors ($\text{\AA}^2$) (min/max/mean) | |
| Protein | 16.77/260.85/112.36 |
| Ligands | 15.72/249.48/97.34 |
| Root mean square deviations |  |
| Bond lengths ($\text{\AA}$) | 0.003 |
| Bond angles (°) | 0.459 |
| **Validation** |  |
| **MolProbity score** | 1.35 |
| Clash score | 2.51 |
| Poor rotamers (%) | 0 |
| **C-beta deviations** | NA |
| **CaBLAM outliers** | 3.74 |
| **Ramachandran plot (%)** |  |
| Favored | 95.48 |
| Allowed | 4.52 |
| Outlier | 0 |

---

**Supplementary Figure 1.**
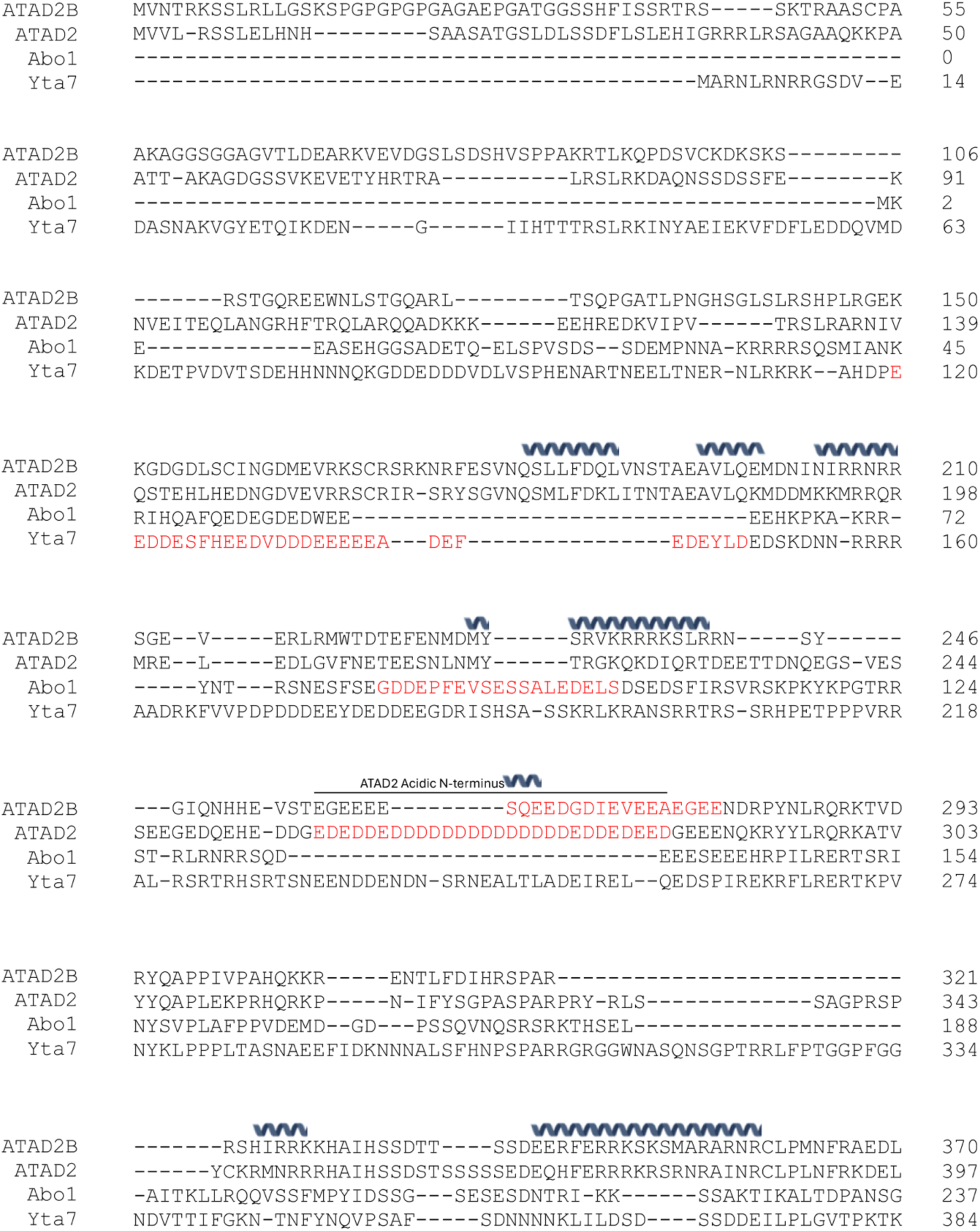
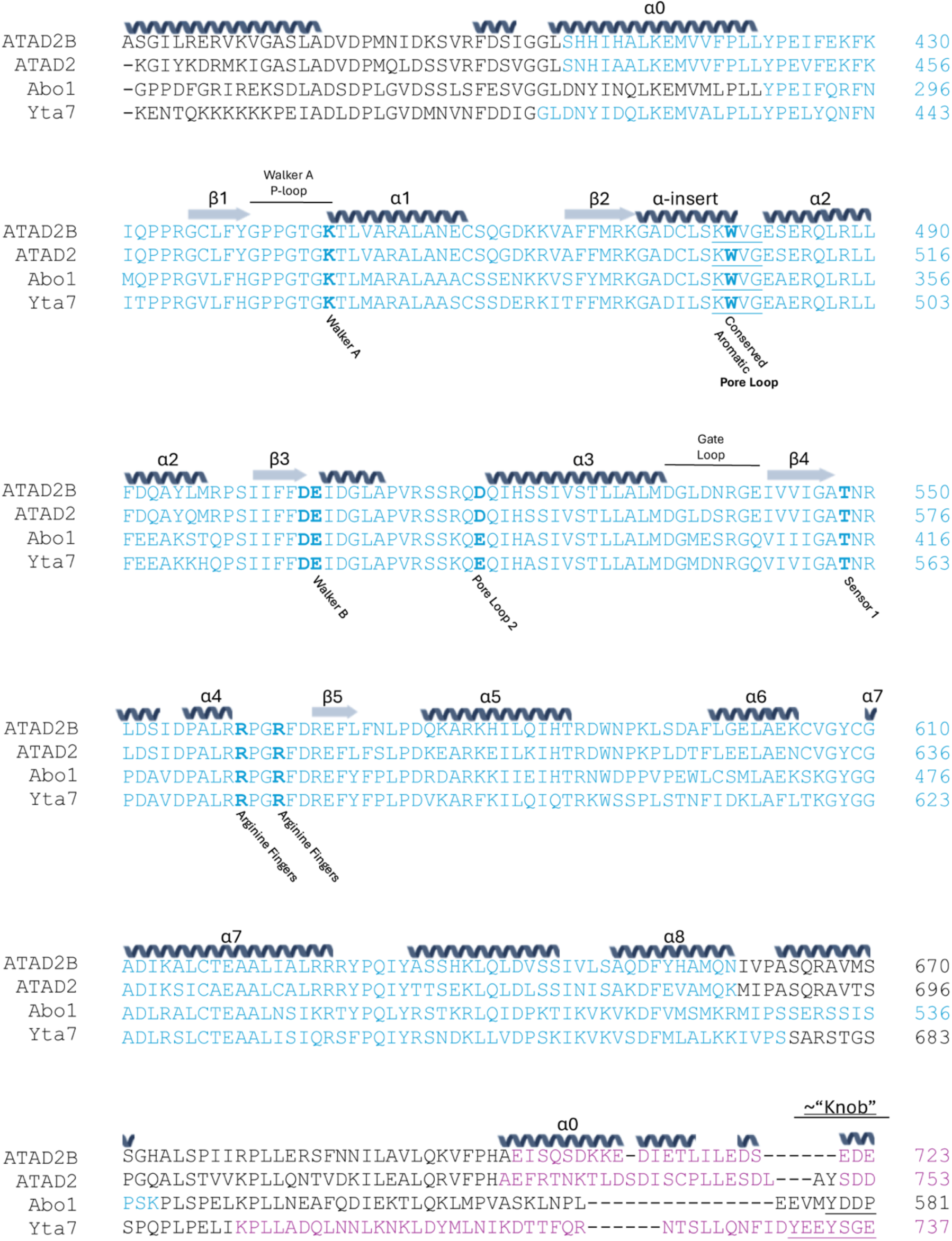
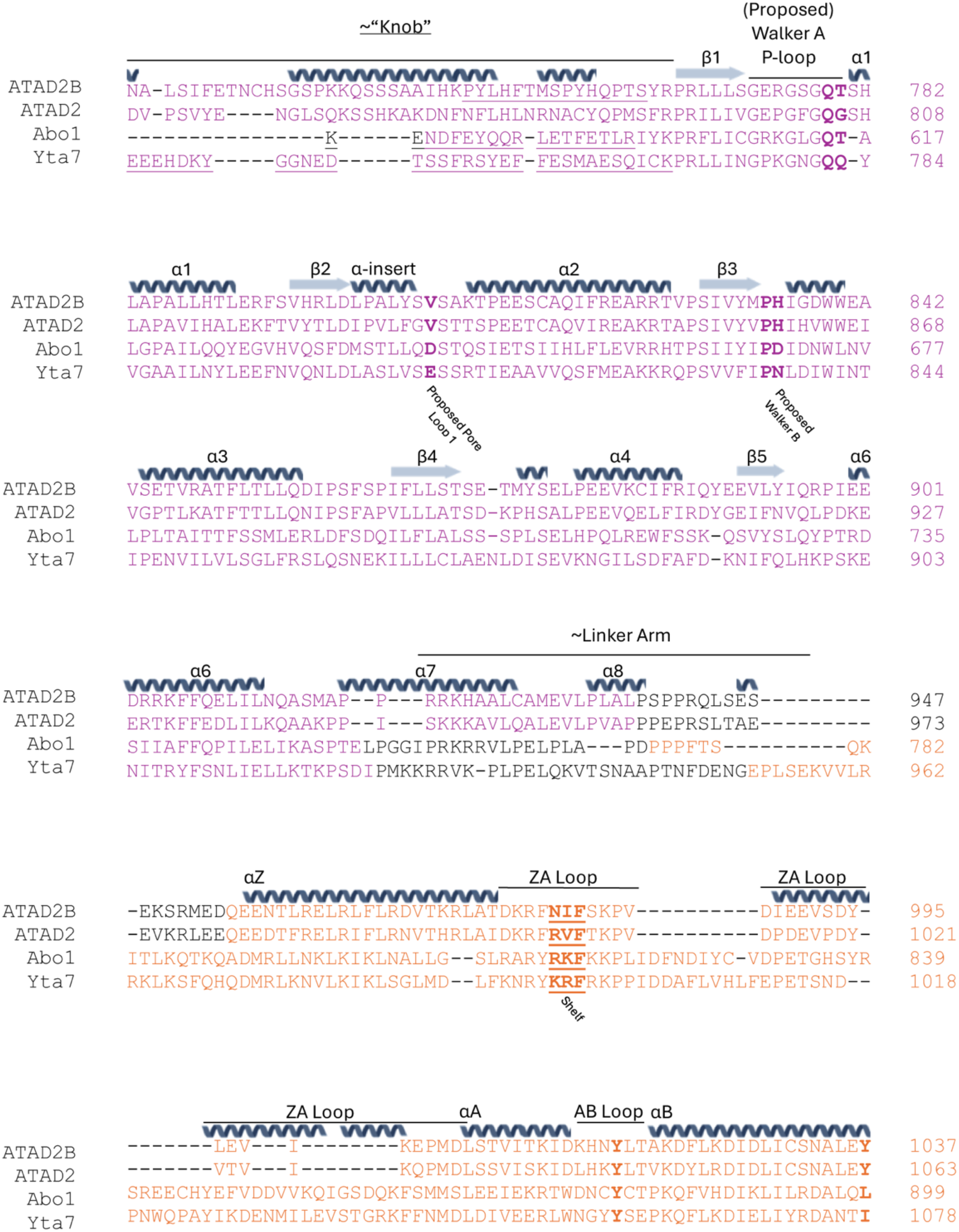
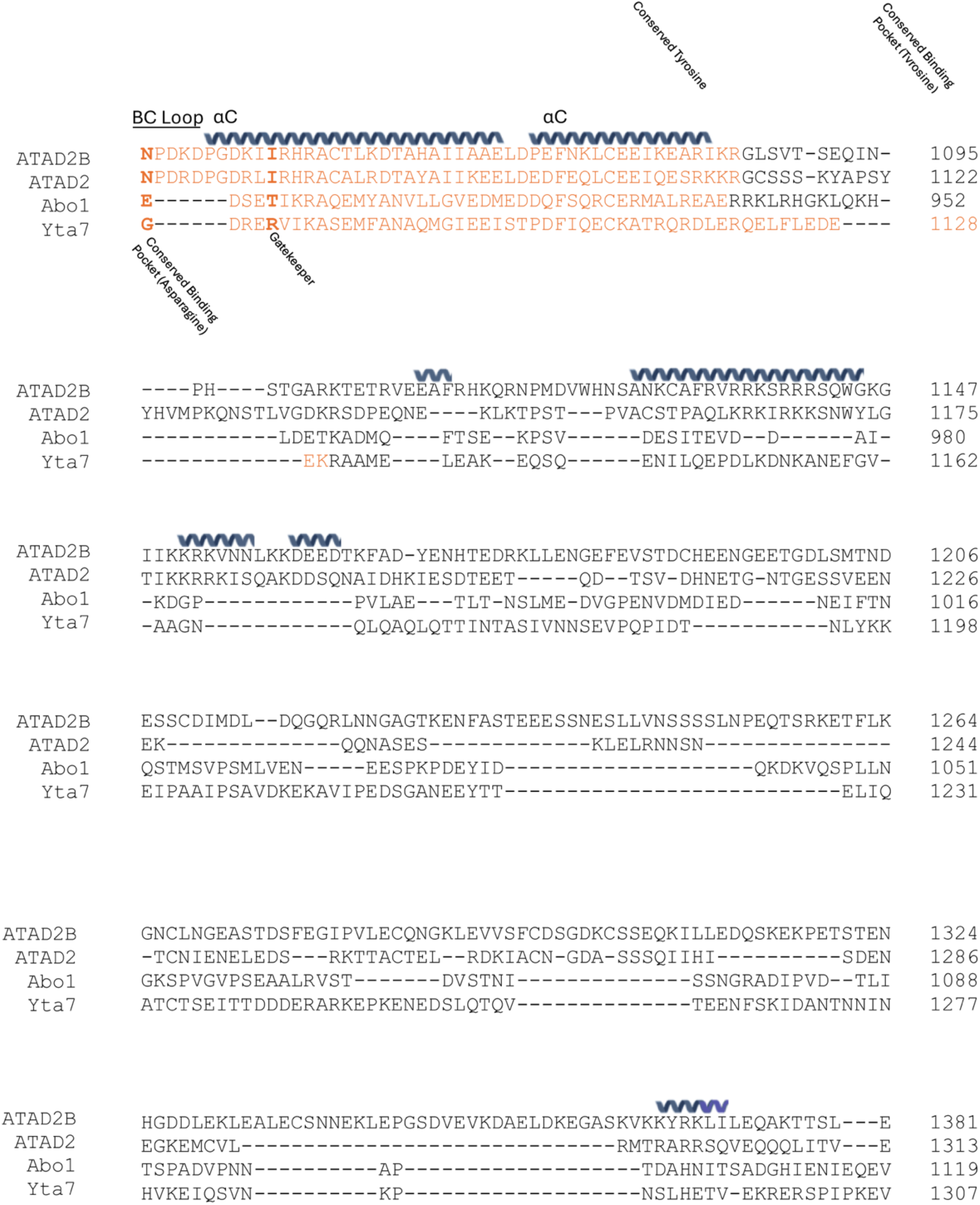
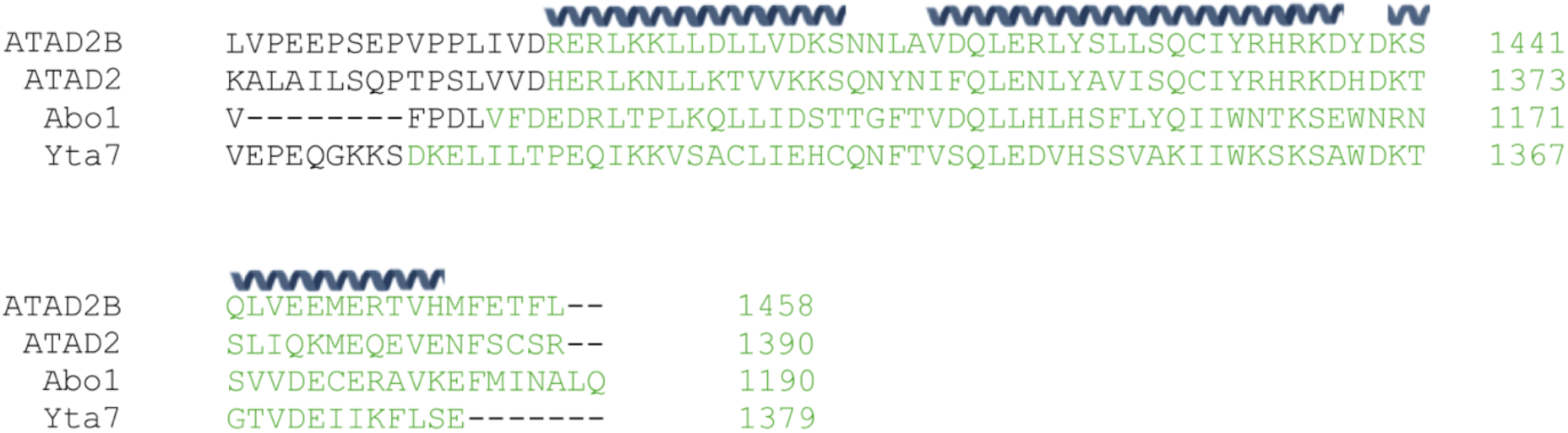
Annotated sequence alignment for the ATAD2-like protein family. Multiple sequence alignment of members of the ATAD2-like protein family with published structures. ATAD2B (UniProt ID: Q9ULI0), ATAD2 (UniProt ID: Q6PL18), Abo1 (UniProt ID: O14114), Yta7 (UniProt ID: P40340) were aligned using the Clustal Omega Multi Sequence Alignment online tool (https://www.ebi.ac.uk/jdispatcher/msa/clustalo). Secondary structure prediction was performed using PROMALS3D^73^. The secondary structure elements are indicated above the sequence, and conserved residues or structural motifs are labeled. The sequence alignment is colored by domain as indicated in Figure 1a where the acidic patch in the N-terminal region is shown in red, the AAA1 domain is blue, the AAA2 domain is purple, the bromodomain is orange, and the C-terminal region is green.

**Supplementary Figure 2.**
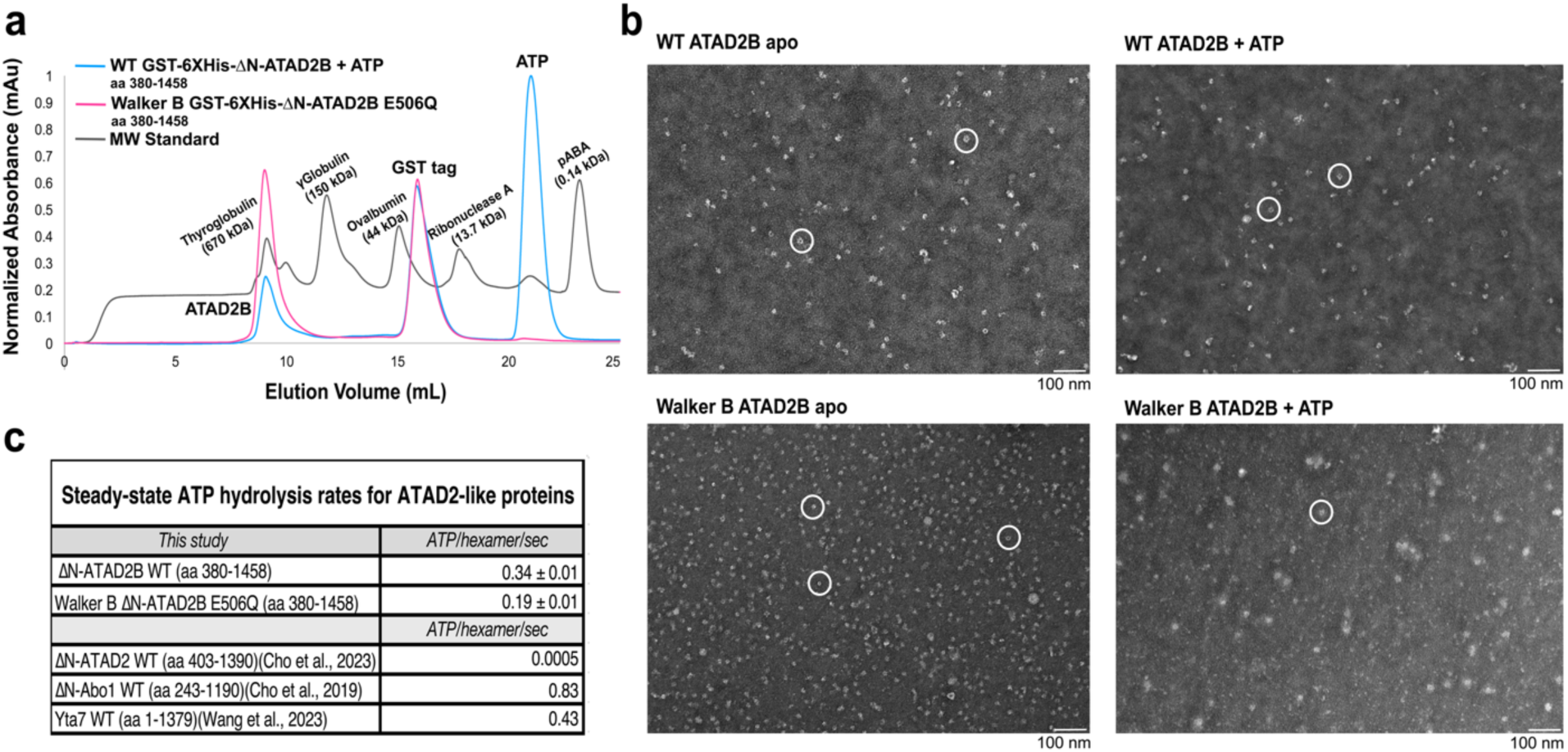
Further investigation into ATAD2B oligomerization. **a)** Size exclusion chromatography (SEC) traces for the wild-type (WT) GST-His6x-ΔN-ATAD2B (residues 380-1458) (blue) and the Walker B (E506Q) GST-His6x-ΔN-ATAD2B Walker B (residues 380-1458) (pink) proteins are overlayed with the molecular weight (MW) standard (grey). The GST-His6x-ΔN-ATAD2B (residues 380-1458) is referred to as the wild-type ATAD2B in this study for clarity. The expected molecular weight for monomeric ATAD2B is ∼150 kDa. **b)** Negative stain electron micrographs captured at 30,000 x on a JEOL 1400 transmission electron microscope (TEM). Hexameric particles are highlighted with a white circle. **c)** Table comparing the steady-state ATP hydrolysis rates of WT ATAD2B and Walker B ATAD2B from this study to the published ATP hydrolysis rates for Abo1^22^, Yta7^25^, and ATAD2^26^. The EnzChek^TM^ phosphate assay kit used in our study was also utilized to measure the rates of ATAD2^26^ and Abo1^22^. The Yta7 results were determined by measuring the phosphate released from ATP hydrolysis^25^.

**Supplementary Figure 3.**
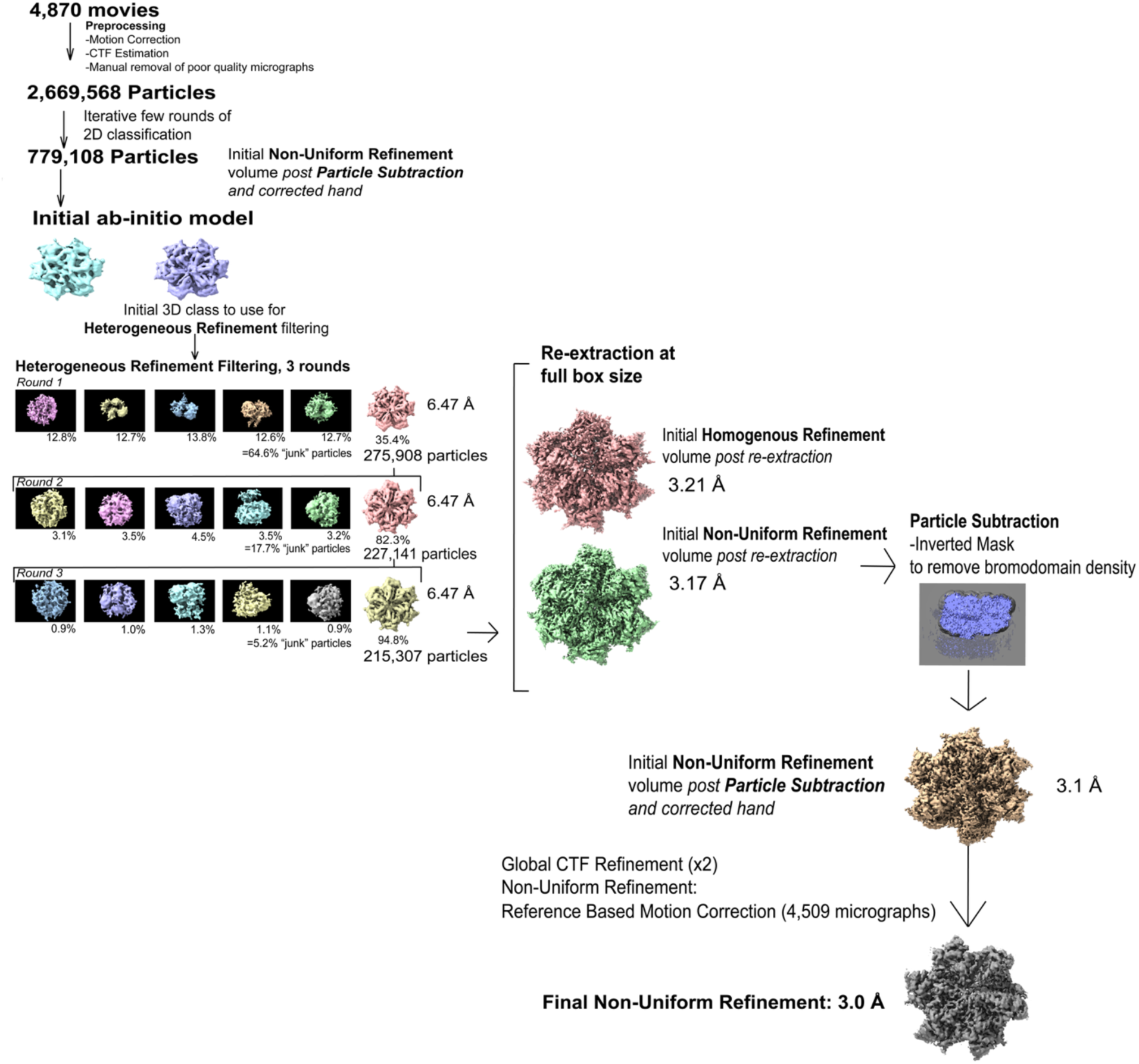
**Overview of the cryo-EM data processing workflow**. The single-particle cryo-EM data processing pipeline used to resolve structural heterogeneity and obtain a high-resolution reconstruction for the ATAD2B Walker B (E506Q) AAA+ ATPase bound to ATPγS and H4K5acK12ac, res 1-24. Following data collection, 4,870 micrographs were preprocessed using patch motion correction, CTF estimation, and particle picking. The resulting 2,669,568 particles were subjected to iterative rounds of two-dimensional (2D) classification, yielding 779,108 particles for downstream analysis. An initial *ab-initio* 3D reconstruction revealed substantial conformational and compositional heterogeneity. Particles were further sorted by multiple rounds of 3D heterogeneous refinement to filter out junk particles. The most homogenous and well resolved class was selected for re-extraction of the particles at the full box size, followed by homogenous refinement, non-uniform refinement, and particle subtraction. Global CTF refinement, non-uniform refinement and reference based motion correction were used to obtain the final 3D reconstruction of ATAD2B with an overall resolution of 3.0 Å. All data processing was carried out with cryoSPARC v.4.5.3^67^.

**Supplementary Figure 4.**
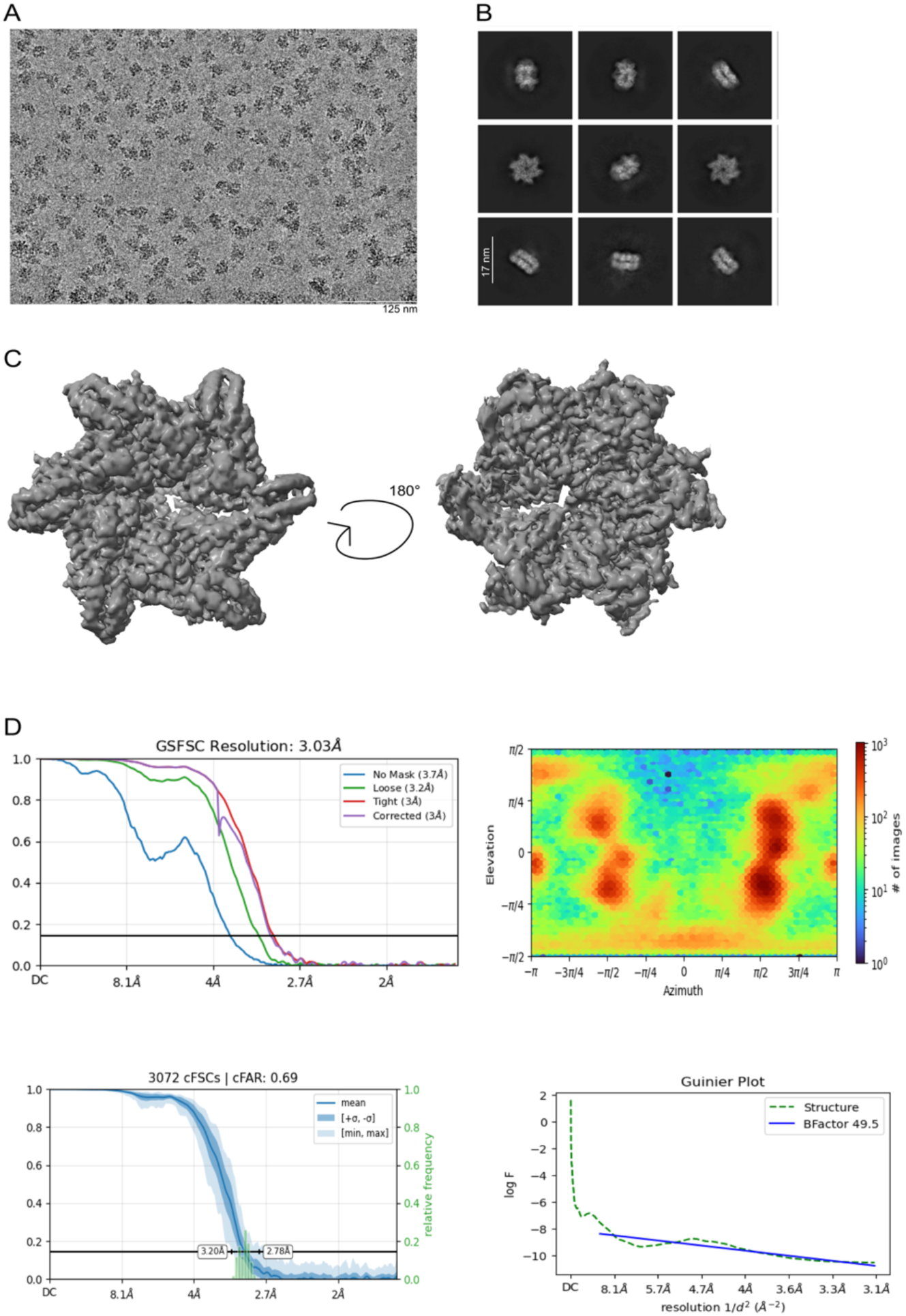
Cryo-EM analysis and resolution assessment of the final ATAD2B Walker B reconstruction. **a)** Representative cryo-electron micrograph showing vitrified ATAD2B Walker B particles embedded in thin ice. Scale bar 125 nm. **b)** Representative 2D class averages from the final particle stack illustrating well-defined particle features in multiple orientations. Scale bar 17 nm. **c)** Final cryo-EM reconstruction of ATAD2B Walker B displaying top and bottom views of the hexameric complex. d) Gold-Standard Fourier Shell Correlation (GS-FSC) curves for the masked, unmasked, and corrected maps of ATAD2B Walker B, with an overall resolution of 3.0 Å at the FSC = 0.143 criterion. **e)** Euler angle distribution plot. **f)** The conical FSC (cFSC) plot reflecting the directional resolution anisotropy. The cFAR value of 0.69 indicates sufficient Fourier sampling. g) Guinier plot comparing the experimental structure factor with and without B-factor sharpening.

**Supplementary Figure 5.**
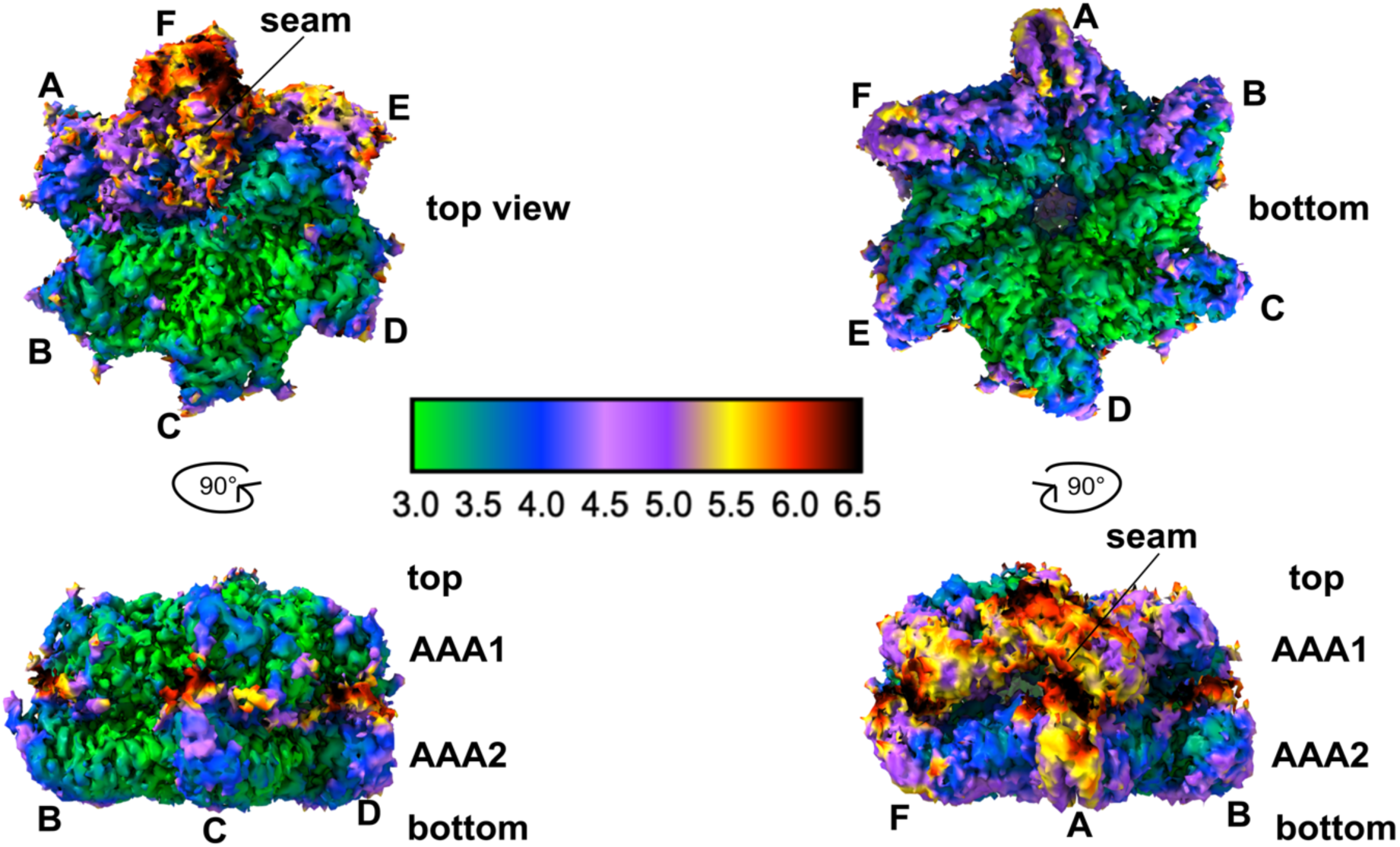
Local resolution assessment of the ATAD2B cryo-EM reconstruction. Top, bottom, interlocking side, bottom, and “seam” side views of the ATAD2B Walker B cryo-EM density map is colored by local resolution in ChimeraX. The local resolution estimation was performed in cryoSPARC v.4.5.3^67^.

**Supplementary Figure 6.**
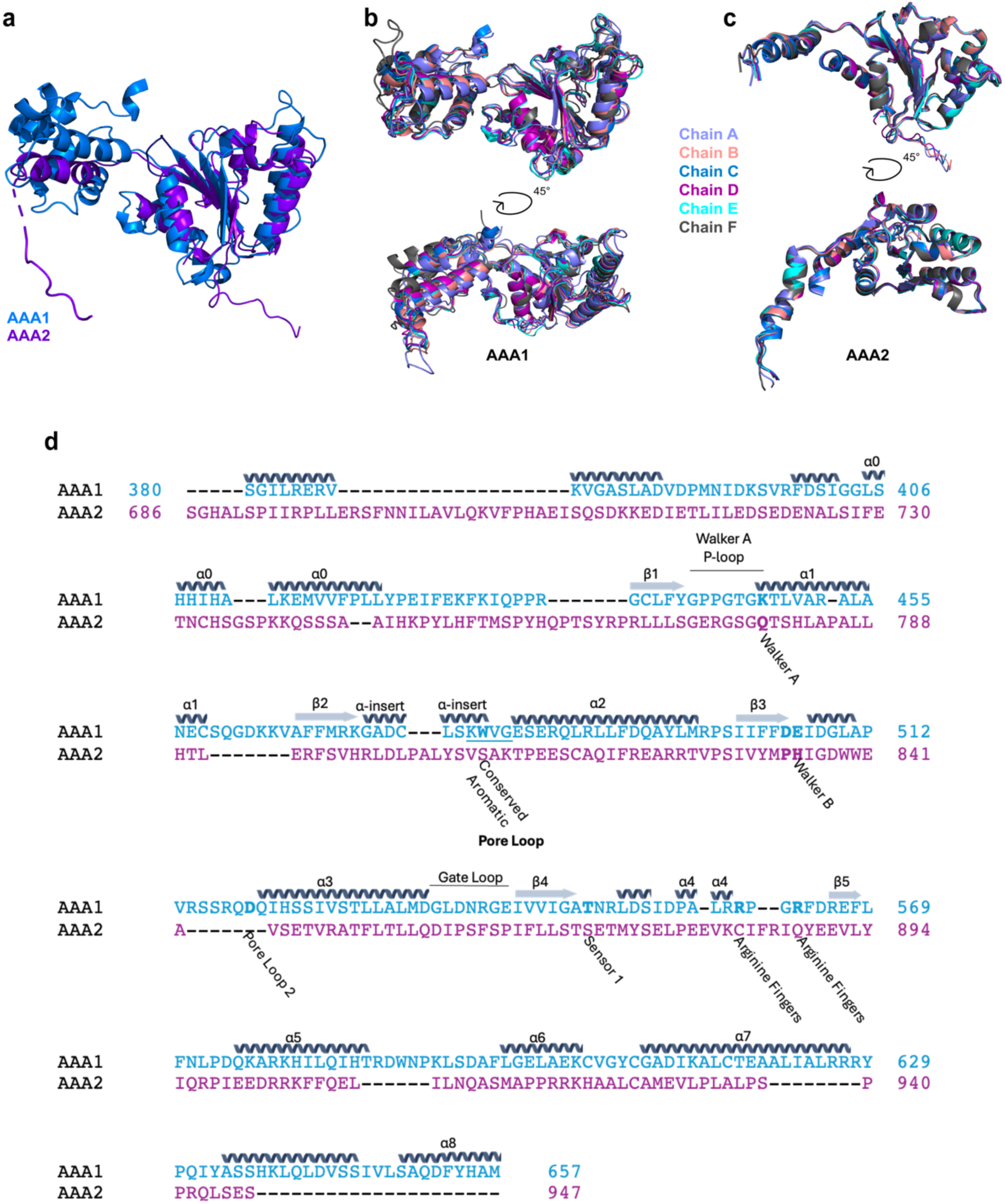
Sequence alignment between the AAA1 and AAA2 domains of ATAD2B. **a)** The AAA2 (purple) and AAA1 (blue) domains from subunit C of the ATAD2B Walker B structure are represented in cartoon and superimposed with each other. **b)** Each of the six AAA1 domains from the ATAD2B Walker B structure are superimposed with each other. Each subunit is colored as indicated. **c)** Each of the six AAA2 domains from the ATAD2B Walker B structure are superimposed with each other. Each subunit is colored as indicated. **d)** Multiple sequence alignment between the AAA1 (blue) and AAA2 (purple) domains of ATAD2B (UniProt ID: Q9ULI0). The domains (numbered as indicated) were aligned using the Clustal Omega Multi Sequence Alignment online tool (https://www.ebi.ac.uk/jdispatcher/msa/clustalo). Secondary structure prediction was performed using PROMALS3D^73^. The secondary structure elements are indicated above the sequence, and the conserved residues or structural motifs are labeled.

**Supplementary Figure 7.**
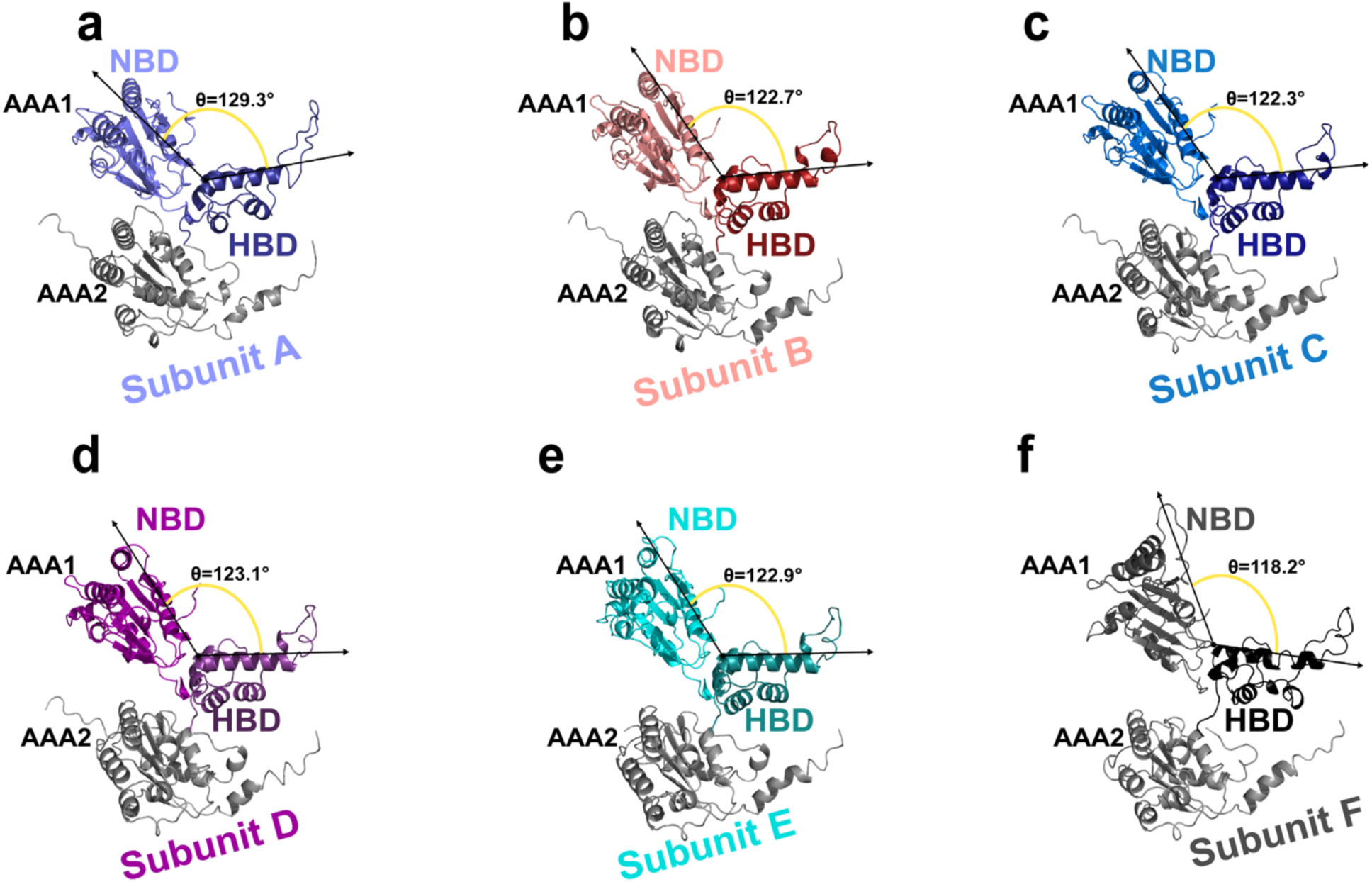
Comparison of each ATAD2B Walker B subunit in the hexameric complex. a-f) Each subunit is displayed in cartoon with the structural features labeled. The AAA1 domains are colored by subunit with subunit A: orchid, subunit B: salmon, subunit C: aqua, subunit D: plum, subunit E: turquoise, and subunit F: iron. In each AAA1 domain the nucleotide binding domain (NBD) is depicted in a lighter shade to the left, and the alpha-helical bundle domain (HBD) is depicted in a slightly darker shade of the same color on the right. All AAA2 domains are colored gray. The orientation of the NBD to the HBD in each AAA1 domain of ATAD2B Walker B was measured using residues 625-611-479, and is represented by two black arrows connected by a yellow curve. The measurement of the angle is displayed next to it and marked with a θ.

**Supplementary Figure 8.**
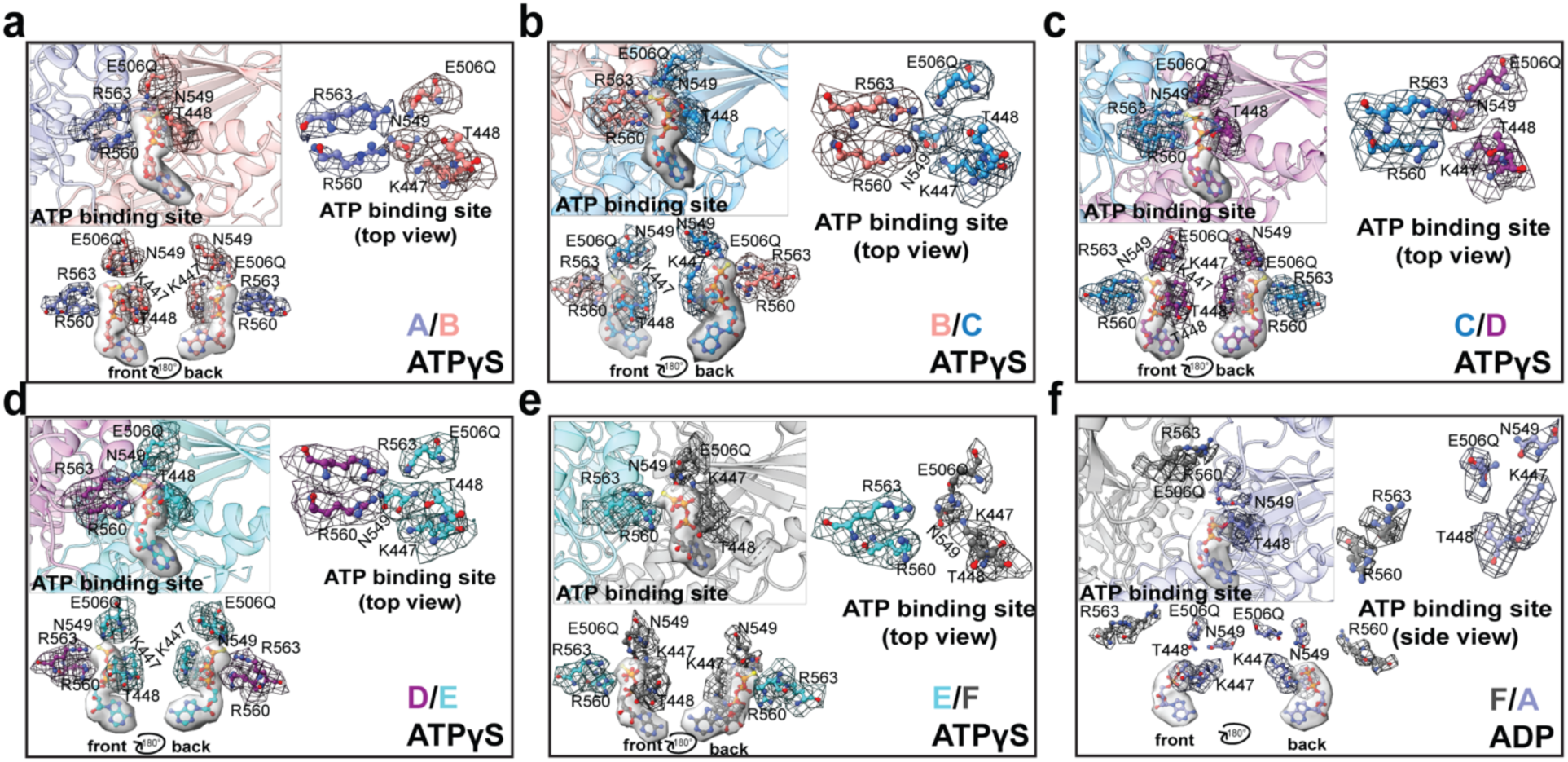
Density in the nucleotide binding site surrounding ATPγS or ADP. a-f) Conserved AAA+ ATPase residues as shown as sticks and include Walker A (K447), T448, Walker B (E506Q), Sensor 1 (N549), and the arginine finger residues (R560 and R563). The cryo-EM density map was contoured at 0.0202 using ChimeraX. For each nucleotide, the density is shown as surface (gray, 41% transparent). For each conserved residue, the density is displayed as mesh and colored the same as its representative subunit according to the following scheme: subunit A: orchid, subunit B: salmon, subunit C: aqua, subunit D: plum, subunit E: turquoise, subunit F: iron. In each panel (a-f), the top left image is a view of the entire ATP binding pocket with the respective subunits shown in cartoon and the conserved residues are shown as sticks. In each panel (a-f), the top right image is a view of the conserved amino acid residues involved in nucleotide coordination without the nucleotide present for better visualization of the density (mesh). In each panel (a-f), the bottom left image is a view of the ATPγS or ADP nucleotide and the conserved amino acid residues involved in nucleotide coordination. Two views are shown, one from the front of the binding site, and another rotated by 180° to show the back for full visualization of each conserved residue.

**Supplementary Figure 9.**
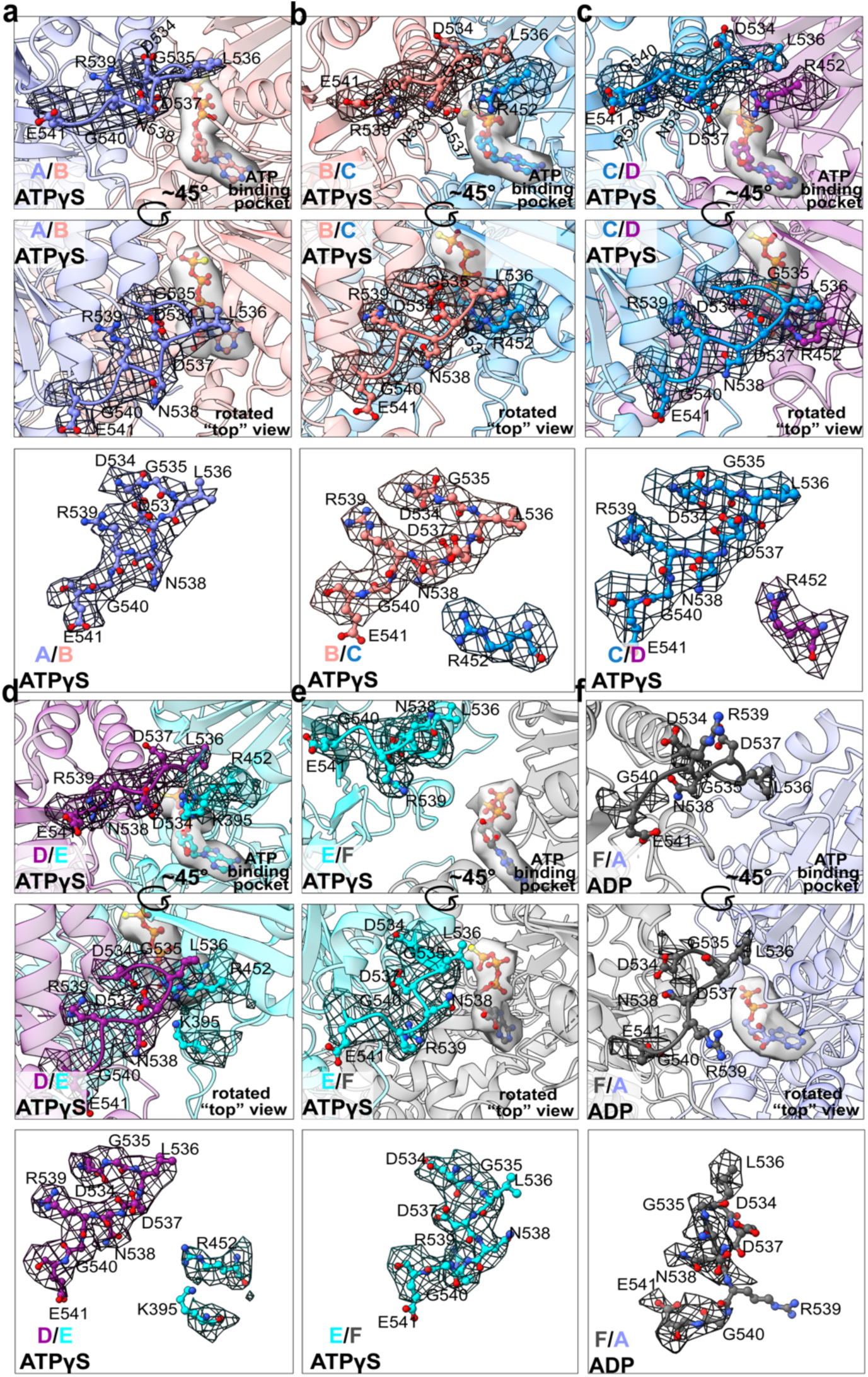
Asymmetric gate loop conformations in the ATAD2B Walker B hexamer. (a-f) Gate loop residues (D534-E541) and interacting residues (R452 and K395) when applicable are shown as sticks. The density map was contoured at 0.0202 in ChimeraX. For each nucleotide, the density is shown as surface (gray, 41% transparent). For each conserved residue, the density is shown as mesh and colored the same as its representative subunit. Subunit A: orchid, subunit B: salmon, subunit C: aqua, subunit D: plum, subunit E: turquoise, subunit F: iron. Three views are shown for a-f, where the top panel is a side view of the gate loop looking into the nucleotide binding pocket. The middle panel is the same gate loop rotated by ∼45° to look into the nucleotide binding pocket. Subunits A-D have “closed” gate loops, and subunits E-F have “open” gate loops. The bottom panel shows the density surrounding the gate loop residues and any nucleotide binding residues when applicable for better visualization of the density.

**Supplementary Figure 10.**
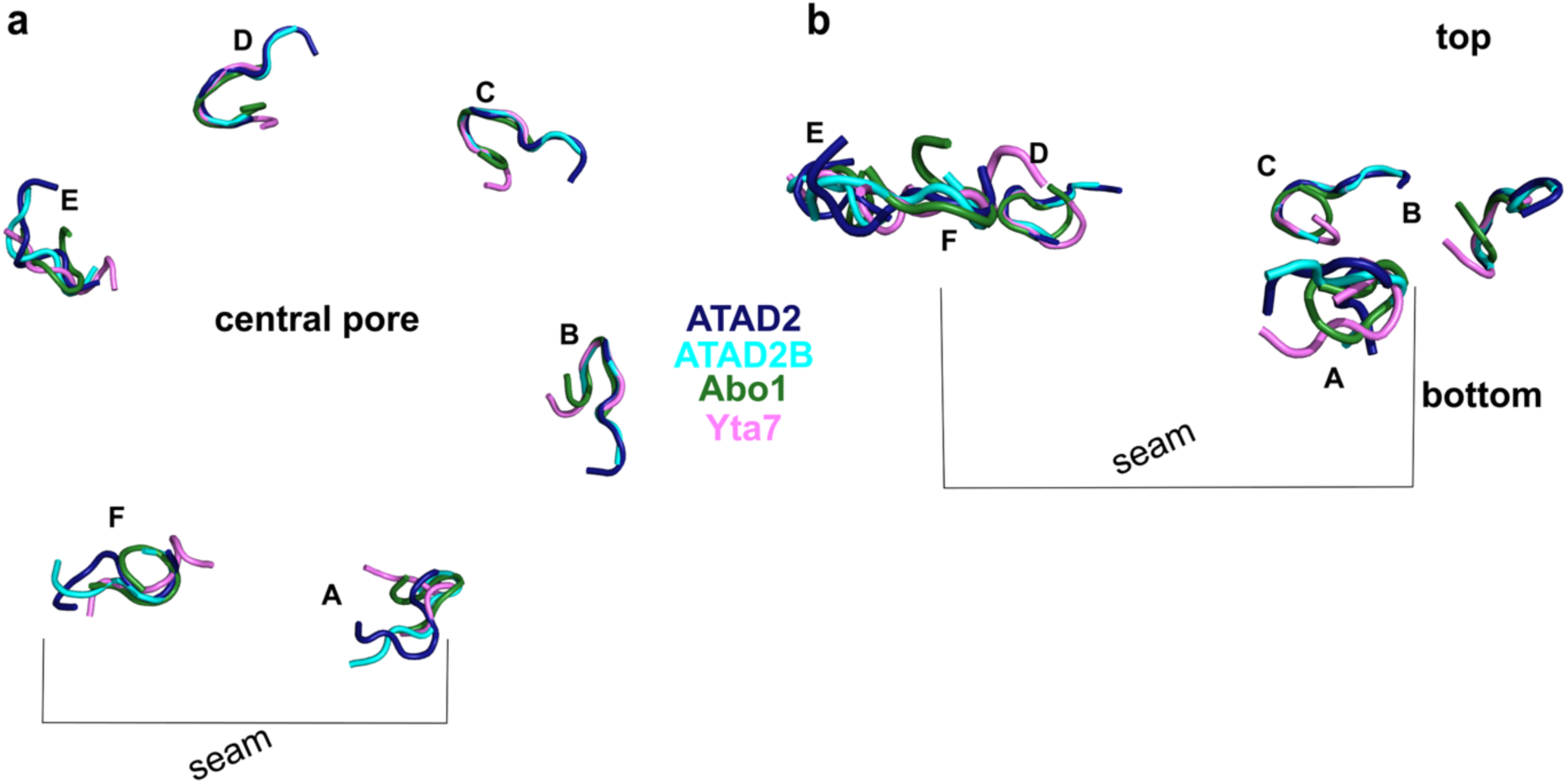
Comparative analysis of the AAA1 domain gate loops across the ATAD2-lke family. a-b) ATAD2B (residues 534-541) was aligned with homologs with solved structures at the gate loops: ATAD2 (PDB ID: 8h3h)(residues 560-568), Abo1 (PDB ID: 6jpu)(residues 398-405), Yta7 (PDB ID: 7uqj)(residues 545-552). These superimpositions are represented as cartoons and colored as indicated: ATAD2B: by subunit, ATAD2: density, Abo1: forest green, Yta7: pink. **a)** Top view of the aligned gate loops from the AAA1 domain. **b)** Side view of the aligned gate loops from the AAA1 domain. Offset between the seam protomers are labeled.

**Supplementary Figure 11.**
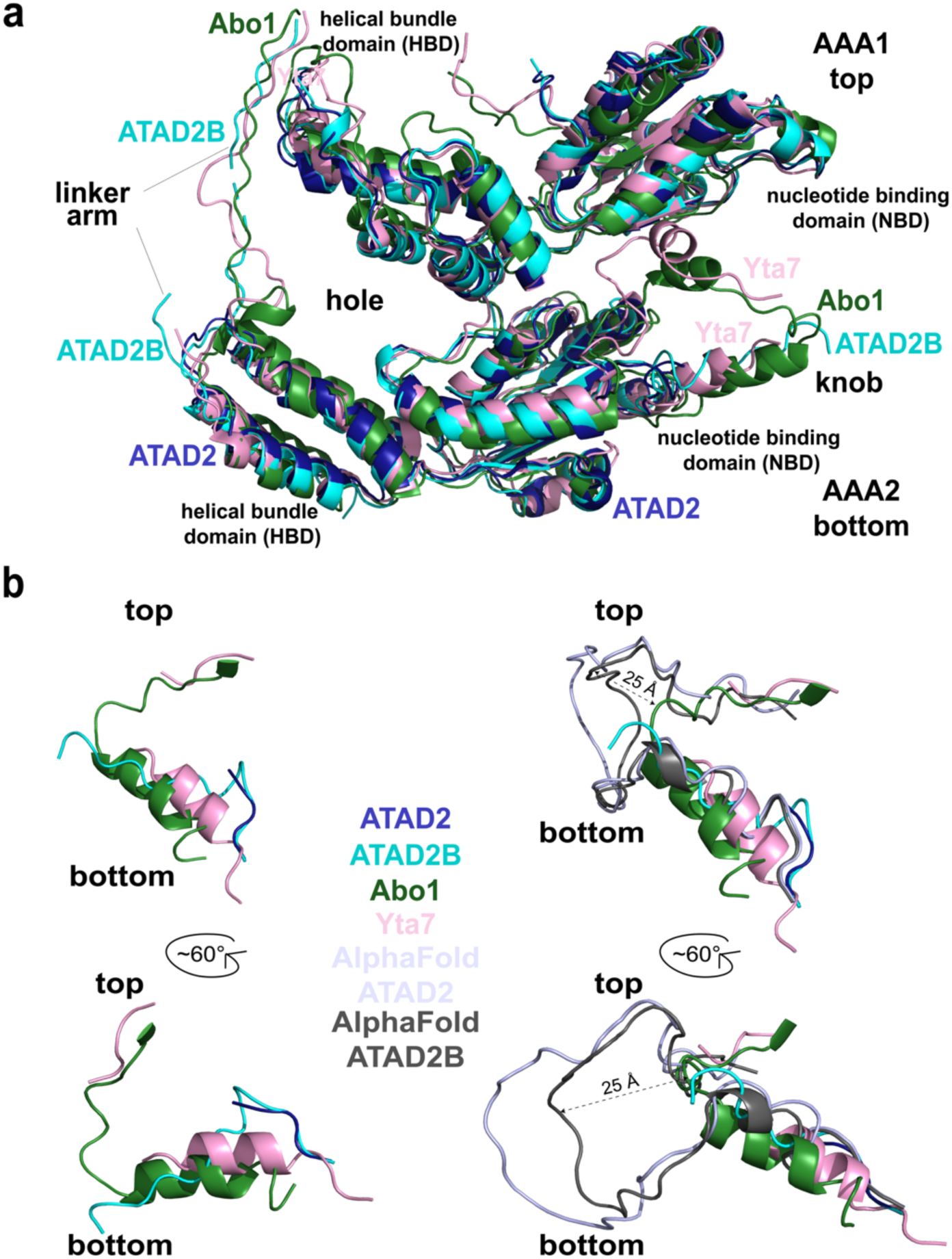
Structural comparison of the knob-hole and linker arm interactions between the ATAD2-like family AAA+ ATPases. a-d) The most well-resolved subunits from the ATAD2B Walker B structure (subunits C and D) were structurally aligned to the corresponding subunits in ATAD2 (PDB ID: 8h3h), Abo1 (PDB ID: 6jpu), Yta7 (PDB ID: 7uqj) using PyMOL^74^. **a)** A surface view of each structure is shown with ATAD2 in purple, ATAD2B in marine, Abo1 in dark blue, and Yta7 in light blue. The knob region from the C subunit of each protein is displayed in cartoon. **b)** The hole is formed by the stacked helical bundle domains of AAA1 and AAA2 along with linker arm from the adjacent subunit (counter clockwise). A cartoon representation of the superimposed D subunits is shown with ATAD2 in hot pink, ATAD2B in purple, Abo1 in violet purple, and Yta7 in light pink. The knob region is also labeled. **c)** The knob-hole region of the ATAD2-like family proteins is depicted. The C and D subunits are shown in surface view with only the knob residues shown in cartoon. The C subunits are colored for each protein with ATAD2 in purple, ATAD2B in marine, Abo1 in blue, and Yta7 in light blue. The knobs extend into the hole formed by subunit D in each protein, which is colored with ATAD2 in hot pink, ATAD2B in purple, Abo1 in violet purple, and Yta7 in light pink. **d)** The left side depicts the knob region of experimentally determined structures. It shows a structural alignment of the knob region from subunit C of each protein with ATAD2 in dark blue, ATAD2B in marine, Abo1 in purple, and Yta7 in green. The right side is the same, but also includes AlphaFold models of ATAD2B (AF-Q9ULI0-F1-v4) in orange, and ATAD2 (AF-Q6PL18-F1-v4) in cyan^44^.

**Supplementary Figure 12.**
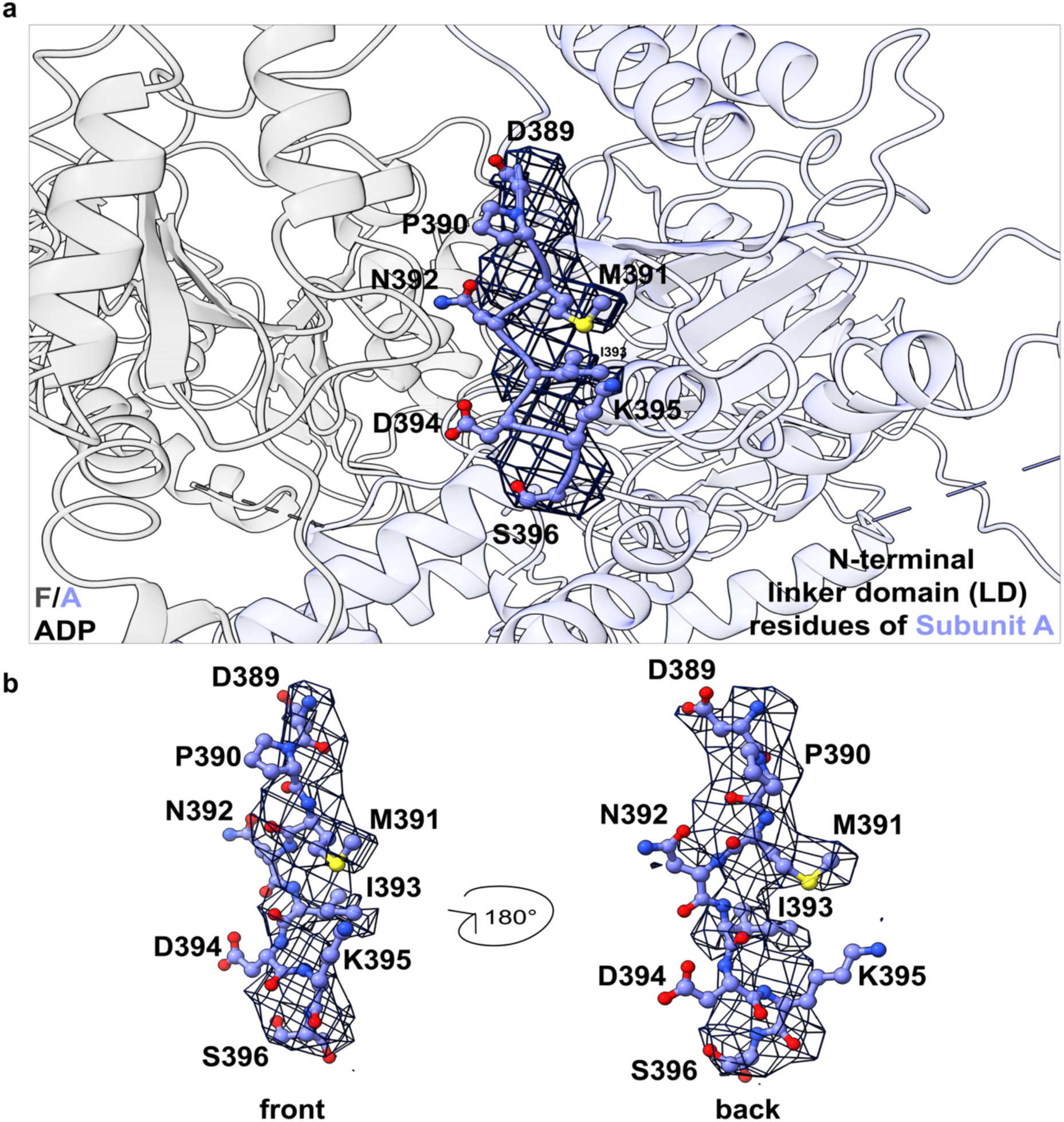
Cryo-EM density of the ATAD2B Walker B linker domain. **a)** The cryo-EM density map of the ATAD2B Walker B-ATPγS-H4K5acK8ac structure was contoured at 0.0202 in ChimeraX. Subunits F (iron) and A (orchid) are represented in cartoon, with residues 389-396 from the linker domain of subunit A shown as sticks. The density map surrounding the linker domain residues is shown as a black mesh around each residue. **b)** Stick representation of the linker domain residues from subunit A with two views rotated by 180° for better density visualization.

**Supplementary Figure 13.**
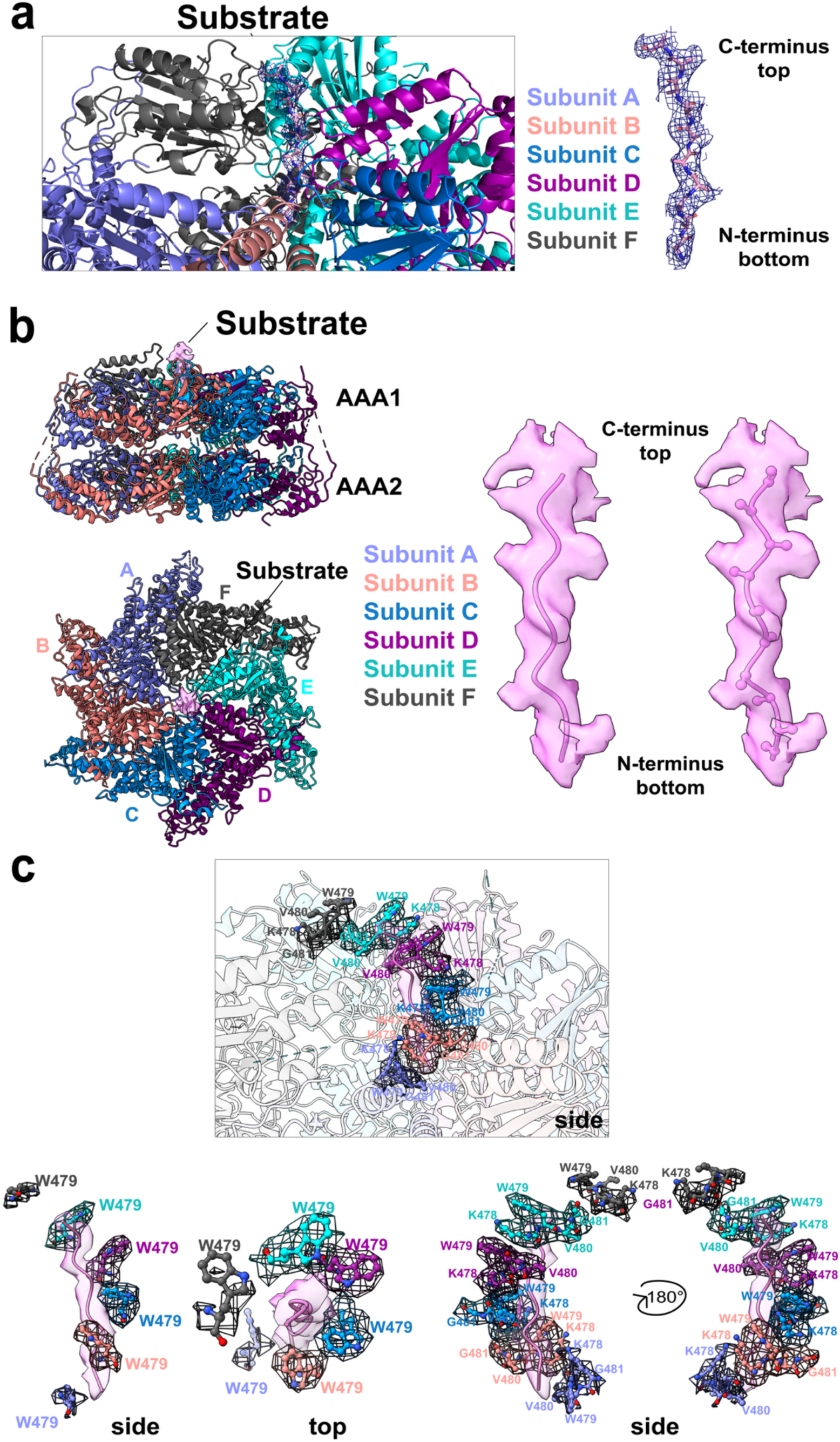
Cryo-EM density of substrate and pore loop residues in ATAD2B Walker B. a-c) Density for subunits A-E was contoured to a level of 0.0202 in ChimeraX, while the density for the more flexible seam subunit F was contoured to a level of 0.0149 for clarity. **a)** Left panel, angled view of the ATAD2B Walker B structure (in cartoon) highlighting density in the pore with each subunit colored as indicated. Right panel, cryo-EM density for the substrate observed in the central pore is shown in blue mesh, and the poly-alanine residues of the modeled substrate are shown in sticks. **b)** Left panel, top and side views of the ATAD2B Walker B structure shown in cartoon. Right panel, left and right images of the cryo-EM density in the central pore shown in pink with cartoon and stick representations of the poly-alanine chain built in the pore. **c)** Top center panel, side view of the tryptophan staircase surrounding the substrate in the central pore region of ATAD2B. The conserved tryptophan residue (W479 in AAA1) is shown in sticks for each subunit in the ATAD2B hexamer and the cryo-EM density surrounding each residue is shown as a black mesh. Bottom panels, show side and top views of the substrate in the pore (cartoon) built into the cryo-EM density (pink). The side view on the right includes the four conserved pore loop residues KWVG (479-481) that contact the substrate with the density in black mesh. The two views of the pore loop residues surrounding the substrate are rotated by 180° for better density visualization.

## Notes

### Competing Interest Statement

The authors have declared no competing interest.

